# Optimisation of the core subset for the APY approximation of genomic relationships

**DOI:** 10.1101/2022.06.06.494931

**Authors:** Ivan Pocrnic, Finn Lindgren, Daniel Tolhurst, William O. Herring, Gregor Gorjanc

## Abstract

**Background:** By entering the era of mega-scale genomics, we are facing many computational issues with standard genomic evaluation models due to their dense data structure and cubic computational complexity. Several scalable approaches have have been proposed to address this challenge, like the Algorithm for Proven and Young (APY). In APY, genotyped animals are partitioned into core and non-core subsets, which induces a sparser inverse of genomic relationship matrix. The partitioning into subsets is often done at random. While APY is a good approximation of the full model, the random partitioning can make results unstable, possibly affecting accuracy or even reranking animals. Here we present a stable optimisation of the core subset by choosing animals with the most informative genotype data.

**Methods:** We derived a novel algorithm for optimising the core subset based on the conditional genomic relationship matrix or the conditional SNP genotype matrix. We compared accuracy of genomic predictions with different core subsets on simulated and real pig data. The core subsets were constructed (1) at random, (2) based on the diagonal of genomic relationship matrix, (3) at random with weights from (2), or (4) based on the novel conditional algorithm. To understand the different core subset constructions, we have visualised population structure of genotyped animals with the linear Principal Component Analysis and the non-linear Uniform Manifold Approximation and Projection.

**Results:** All core subset constructions performed equally well when the number of core animals captured most of variation in genomic relationships, both in simulated and real data. When the number of core animals was not optimal, there was substantial variability in results with the random construction and no variability with the conditional construction. Visualisation of population structure and chosen core animals showed that the conditional construction spreads core animals across the whole domain of genotyped animals in a repeatable manner.

**Conclusions:** Our results confirm that the size of the core subset in APY is critical. The results further show that the core subset can be optimised with the conditional algorithm that achieves a good and repeatable spread of core animals across the domain of genotyped animals.

## Background

Estimating breeding values is a key operation in identifying the best animals. Estimated breeding values (EBVs) are usually obtained with the best linear unbiased prediction (BLUP), where genetic covariances between animals enable sharing of information between relatives and improve accuracy of estimation. Traditionally pedigree has been used to construct a matrix of expected genetic covariances **A** (the pedigree relationship matrix). With the advent of genome-wide single nucleotide polymorphism (SNP) markers, realised genetic covariance matrix based on SNP genotypes **G** (the genomic relationship matrix) replaced the pedigree-based **A**. This change gave us the so called genomic BLUP (GBLUP) and genomic EBVs (GEBVs). An essential part of estimating GEBVs via Henderson’s mixed model equations is the inversion of **G** [1]. Matrix inversion has a cubic computational cost and becomes a limiting step with more than ∼150,000 genotyped animals [2]. A similar computational bottleneck was inverting the **A** with the pedigree BLUP. This bottleneck was removed with a recursive algorithm of setting up the sparse inverse of **A** directly from a pedigree [3, 4]. Unfortunately, such an algorithm is not available for **G** because genome-wide SNP genotypes do not have a recursive data structure and the inverse of **G** is dense. This is an issue because the number of genotyped animals in increasing rapidly in many populations. The most remarkable case is in the US Holstein dairy cattle, where already 5 million animals have been genotyped (The Council of Dairy Cattle Breeding, https://www.uscdcb.com).

Several approaches have been proposed to solve large GBLUPs. For example, reparameterizing the model with marker effects [5, 6], inducing sparsity in the inverse through the approximation of genomic relationships [7], leveraging matrix inversion lemma [8], or using dimensionality reduction [8, 9]. The Algorithm for Proven and Young (APY) induces sparsity in the inverse through the approximation of genomic relationships by leveraging the limited dimensionality of genomic information in populations with a small effective population size [10]. This approximation is achieved by splitting genotyped animals into core and non-core subsets. The core animals are assumed to be all dependent on each other and hence we construct the full inverse for these animals. The non-core animals are assumed to be conditionally independent given the core animals, hence the inverse for these animals is diagonal, while the part of the inverse between the core and non-core animals is dense. With APY, direct inversion is needed only for the core subset. Effectively, GEBVs of the non-core animals are modelled as a function of the GEBVs of the core animals. During model fitting, phenotype information “flows” between both subsets of animals, hence both subsets contribute to the estimation. While APY is an approximation, it has been shown to be accurate and scalable for various populations of dairy and beef cattle [11, 12], pigs [13, 14], and sheep [15, 16].

There are two crucial decisions in APY. First, how many animals should form the core subset. Second, which animals should form the core subset. Empirical testing suggests that the optimal number of core animals is connected to the effective population size and the dimensionality of genomic information [10]. As such, the optimal number of core animals can be gauged by the number of eigenvectors that explain a large percentage of variation in SNP genotypes (or uquivalently in **G**) [17]. For example, by setting the number of core animals to the number of eigenvectors that captured more than 98% of the variation in **G**, the GEBVs were comparable to those obtained with the full **G**. Empirical testing also suggests that a random choice of core animals gives satisfactory accuracy of GEBVs [18]. However, random choice is not desired in routine genetic evaluations because it could result in small changes in GEBVs, possibly reranking animals, even with the same data. A recent study suggested that such changes are a natural part of genetic evaluations when the data is updated [19]. Still, genetic evaluations should return the same GEBVs with the same data. An extensive study on size and definitions of core subset in pig dataset [20] showed that the definition of core subset matters for small core sizes. Therefore, there is a need for methods that construct an optimal core subset that maximises accuracy of GEBVs and ensures stable results between different model runs.

The aim of this paper is to present and evaluate alternative core subset constructions for APY. First, we propose a novel algorithm for optimising the core subset based on the conditional genomic relationship matrix or the conditional SNP genoypes matrix. We termed this method as the conditional core subset algorithm, or simply conditional algorithm. Second, we compared the conditional core subset construction with other core subset constructions on a simulated cattle data and a real pig data.

## Methods

We analysed how different core subset sizes and different core subset constructions affect the accuracy of GEBVs with APY. Specifically, we compared core subset construction (1) at random; (2) based on the diagonal of the genomic relationship matrix; (3) at random with weights from (2); or (4) based on the conditional algorithm. We compared the different core subset constructions with the number of core animals that captured 10, 30, 50, 70, 90, 95, 98, and 99% of the variation in **G**. To demonstrate the functionality of different core subset constructions we used a small simulated cattle data. To demonstrate their practical utility we used a large real pig data. In the following, we first describe the simulated and real data and the models we fitted to each dataset. We then describe different core subset constructions for APY, including the derivation of the conditional algorithm. Finally we describe the validation of GEBVs and data visualisation.

### Simulated cattle data

We simulated a simple breeding programme using the `AlphaSimR R` package [21]. The simulated genome consisted of 10 chromosomes with 1100 segregating sites per chromosome (11,000 total), from which we randomly assigned 100 sites (1000 total) as additive quantitative trait loci (QTL), and 1000 sites (10,000 total) as SNP markers. There was no overlap between QTLs and SNPs. The historical effective population size followed cattle estimates [22], with mutation rate of 2.5 × 10^−8^, and recombination rate of 1.0 × 10^−8^. Phenotypes were assumed to have heritability of 0.30. From the initial founder population of 3000 animals, we randomly selected 1500 females and 50 males to serve as parents of the first generation. In each succeeding generation, 3000 animals with equal female to male sex ratio were obtained from mating 1500 females and 50 males, with a fixed number of two offspring per female. Females were replaced at 50% rate, meaning that the 750 best young females out of 1500 and the 750 best old females out of 1500 were dams of the next generation. This dam selection was based on their phenotype value. On the male side, the best 45 young males out of 1500 and the best 5 old males out of 45 were sires of the next generation. This sire selection was based on their true genetic value (mimicking accurate selection). SNP genotypes were collected for all animals in generations 15-20, resulting in 15,000 genotyped and phenotyped animals in our study. The 3000 genotyped animals from generation 20 served as a validation subset.

#### Simulated cattle data analysis

We analysed the simulated phenotype and SNP genotype data with GBLUP using:

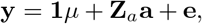

where **y** is a vector of phenotypes, **1** is a vector of 1’s, *µ* is the overall mean, **Z**_*a*_ is a design matrix connecting phenotypes to animal breeding values **a**, and **e** is a vector of residuals. We assumed that 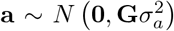 and 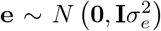, where **G** is the genomic relationship matrix, 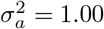 is variance of breeding values, and 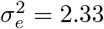 is variance of residuals.

### Real pig data

Phenotypic data for a moderately heritable trait measured from 1999 to 2021 on 42,868 pigs (33,544 purebred - L1, 114 crossbred - F1, and 9,210 backcross - BC1 and BC2) were provided by PIC (a Genus company, Hendersonville, TN, USA). SNP genotypes were available for 42,707 SNP markers after quality control. The number of genotyped pigs was 49,788 pigs (37,598 purebred - L1, 486 crossbred - F1, and 11,704 backcross - BC1 and BC2). The validation subset included 478 phenotyped and genotyped youngest animals (L1 and BC2) born in 2021, with their phenotypes removed from the analysis.

#### Real pig data analysis

We analysed the real phenotype and SNP genotype data with GBLUP using:

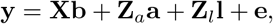

where **y** is a vector of phenotypes, **X** is a design matrix connecting phenotype to contemporary group fixed effects **b, Z**_*a*_ is a design matrix connecting phenotypes to animal breeding values **a, Z**_*l*_ is a design matrix connecting phenotypes to shared litter effect **l**, and **e** is a vector of residuals. We assumed that 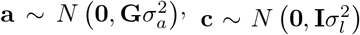,and 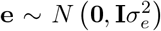 where **G** is the genomic relationship matrix, 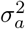 is variance of breeding values, 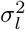 is variance of litter effects, and 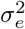 is variance of residuals.

### Genomic relationship matrix and APY inverse

For both simulated and real data, genomic relationship matrix was calculated as 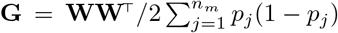, where **W** is a centred matrix of SNP genotypes (coded as 0 for reference homozygote, 1 for heterozygote, and 2 for alternative homozygote), *p*_*j*_ is the frequency of alternative allele for SNP marker *j*, and *n*_*m*_ is the number of SNP markers [23]. To ensure **G** is positive definite, **G** was blended with 0.01**I**. The final **G** was either inverted directly or with APY.

The APY partitions **G** into:

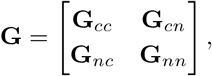

where *c* and *n* correspond to the core and non-core subsets. The APY inverse [7] is then:

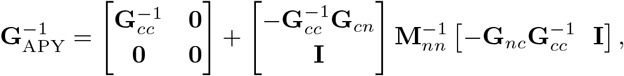

where 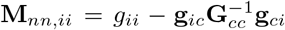 in which *g*_*ii*_ is the *i*^*th*^ diagonal element of **G**_*nn*_ and **g**_*ci*_ = **g**_*ic*_ is the *i*^*th*^ column of **G**_*cn*_ (or the *i*^*th*^ row of **G**_*nc*_). This formulation induces a sparse inverse that approximates the full inverse. Here we only need the direct inverse of **G**_*cc*_ and the diagonal matrix **M**_*nn*_.

We defined the core and non-core subsets by splitting the genotyped animals. The number of core animals was the same across all evaluated core subset constructions. We gauged the number of core animals with the number of eigenvectors that captured 10, 30, 50, 70, 90, 95, 98, and 99% of variation in **G** [17]. The remaining genotyped animals then formed the non-core subset.

Instead of eigenvalue decomposition of **G**(*n* × *n*) we used an equivalent singular value decomposition (SVD) of **W**(*n* × *n*_*m*_), because SVD has a 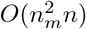 computational cost, while eigenvalue decomposition has a *O*(*n*^3^) computational cost. The SVD is given by **W** = **UDV**^⊤^, where **D** is a diagonal matrix of singular values that correspond to the square root of the non-zero eigenvalues of **W**^⊤^**W** and **WW**^⊤^. The columns of **U** are left singular vectors that correspond to the eigenvectors of **WW**^⊤^, such that **U**^⊤^**U** = **I**. The columns of **V** are right singular vectors that correspond to the eigenvectors of **W**^⊤^**W**, such that **V**^⊤^**V** = **I**. The eigenvalue decomposition of **G** is then obtained as **G** = **WW**^⊤^*/v* = **UD**^2^**U**^⊤^*/v*, where 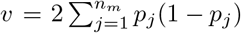. The SVD of **W** was computed with the base `R` function `svd` [24] for the simulated data and a `Fortran` program using `LAPACK` subroutine `DGESVD` [25] for the real pig data.

### Core subset constructions

A guiding principle in APY is to construct a core subset in such a way that APY inverse approximates the full inverse **G**^−1^ well. This can be achieved by a large enough core subset. Further, for a given size of a core subset, we want to cover as much variation in SNP genotype matrix (and hence in **G**) as possible. To achieve this, we used four different core subset constructions. The first construction (random), randomly sampled genotyped animals to form the core. This construction is the most common in current APY applications and has hence served as a baseline. The second construction (diagonal), chose core animals based on diagonal elements in **G** [15]. The principle behind this construction is that animals with large diagonal elements in **G** deviate substantially from the centroid of SNP genotypes and therefore sample variation in SNP genotypes well. An alternative view of this principle is to recognise that the trace of a matrix is equal to the sum of its eigenvalues. Hence, selecting animals with the largest diagonal elements in **G** maximises the amount of captured variation. However, this construction can oversample individuals from the most inbred families, while we would like to cover as many families as possible for a given core subset size. The third construction (weighted), was a combination of the first two approaches, where the core animals were randomly sampled, but with a weight based on the corresponding diagonal element in **G**. The weighted construction attempted to alleviate the potential oversampling issue with the diagonal construction. We used the `R` function `sample n` [26] with option `weight` for weighted sampling. For the random approaches (random and weighted), sampling was replicated five times to manifest potential variability in GEBVs with a random core subset, and in the results we present the mean and 95% quantiles across the replicates. The forth construction (conditional), was based on a conditional algorithm inspired by sequential sampling design, which we describe in the following.

The principle behind the conditional algorithm is to spread core animals across the domain of genotyped animals or equivalently the domain of collected SNP genotype data. This is achieved by choosing animals far away from each other in covariance sense. The algorithm initiates the core subset with the animal that deviates the most from the centroid of SNP genotypes. Then, it sequentially finds animals that deviate the most from the core animal(s). By growing the core subset the algorithm optimally samples the domain.

Before we introduce the algorithm, we will show how to find animals that are far away from each other in covariance sense. We start with the joint covariance matrix **C** = **WW**^⊤^, where the variance for animal *i* is the squared norm of the *i*-th row of **W**, or equally the *i*-th diagonal element of **WW**^⊤^. For simplicity we omitted the scaling constant, so **G** = **C***/v*. Let **w** denote the vector corresponding to the *i*-th row of **W** (SNP genotypes of the animal *i*). Let **e** denote a “selector” vector of 0’s and a single 1 at the position *i*, so that **w** = **e**^⊤^**W, C**_*i,i*_ = **e**^⊤^**Ce**, and **C**_*i*,−*i*_ = **e**^⊤^**C**. To find animals that are far away from each other in covariance sense we use conditional covariance matrix **C**_*cond*_ where we condition on core animals. Specifically, large diagonals of **C**_*cond*_ indicate animals whose SNP genotypes are not well represented by SNP genotypes of core animals. We implemented the algorithm by expanding the core subset one animal at a time. Hence, we require conditional covariance matrix given the animal *i*, which is:

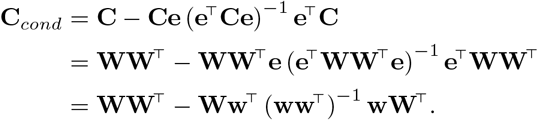

Since **ww**^⊤^ is the squared norm (||**w**||2) of the row vector **w**, we introduce a normalised vector **a**, where 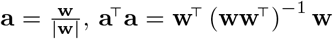, and **aa**^⊤^ = 1. By using **a**, we can shorten the expression for conditional covariance to:

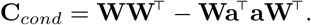

By adding and subtracting **Wa**^⊤^**aW**^⊤^ we can factorise the conditional covariance as:

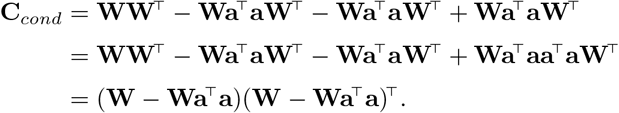

The above factorised expression shows that we can calculate conditional SNP genotype matrix (conditional on the SNP genotypes of the animal *i*) as:

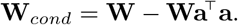

Therefore, conditional covariance matrix is:

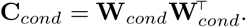

The conditional algorithm (Algorithm 1) starts with a desired size of the core subset *n*_*c*_, vector to store core animals, and the joint covariance matrix **C**_0_ = **WW**^⊤^. The algorithm iterates for *n*_*c*_ rounds by sequentially choosing core animals based on their conditional variance. In the *i*-th round, it expands the core subset with an animal that has the largest conditional variance in **C**_*i*−1_, that is, conditional on the (*i* − 1) previously chosen core animals. Then it updates **C**_*i*−1_ to **C**_*i*_ by conditioning on the currently chosen core animal. In the next round, the updated conditional covariance matrix **C**_*i*_ is used to choose the next core animal, and so on. Note that if the covariance matrix is not updated, we obtain the diagonal construction.

#### Algorithm 1

Core subset optimisation using conditional covariance matrix **C**

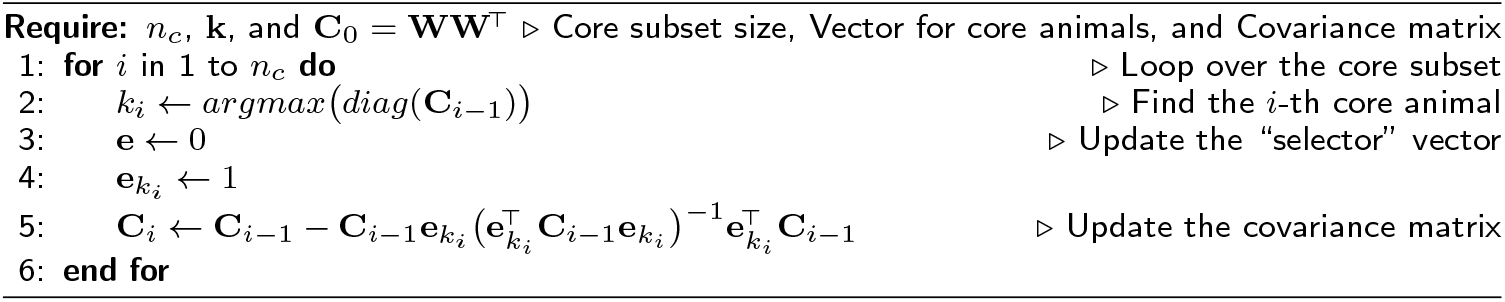

Forming and repeatedly updating a large covariance matrix can be consuming in terms of compute and storage. The conditional algorithm can work with the SNP genotype matrix instead of the covariance matrix by using the expression for conditional SNP genotype matrix (**W**_*cond*_) (Algorithm 2). In line 2 of this algorithm, we show expression 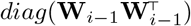, which forms the covariance matrix and extracts its diagonal. We can speedup this step by calculating only the diagonal (conditional variance) by squaring and summing every row of **W**_*i*−1_.

#### Algorithm 2

Core subset optimisation using conditional SNP genotype matrix **W**

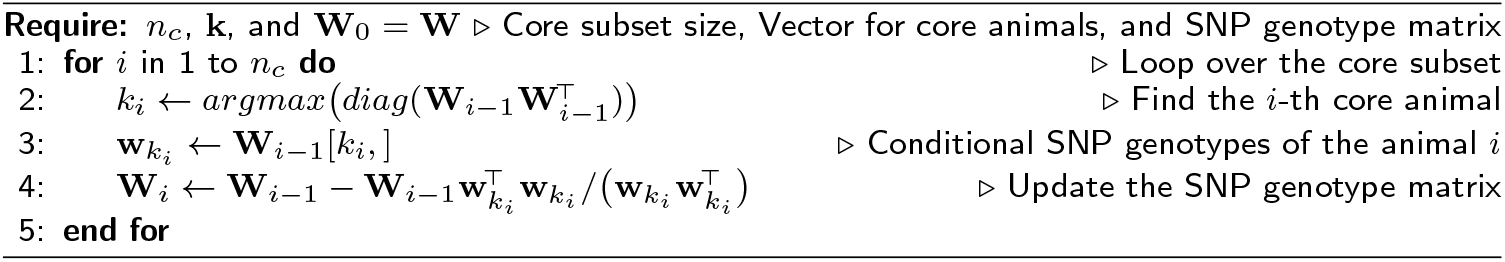

The above presented algorithm does not guarantee a globally optimal core subset. Namely, a different starting core animal will lead sequential updates to a different core subset. Achieving global optimality is demanding. To partially address the optimality issue, we have extended the algorithm to choose core animals that sequentially minimise conditional variances of other animals. That is, choosing a core animal that captures as much variation in SNP genotypes of other animals as possible. This principle follows closely the principle of APY approximation. We have derived two versions of the extension by sequentially minimising maximum or average conditional variance of other animals. To implement the extended algorithm, we have to replace the line 2 of Algorithm 2 with its line 4 and run this updated line for every animal. Unfortunately, the addition of this inner loop makes the extended algorithm impractically slow and we have not tested it in our further work.

### Analyses

We compared different core subset constructions with correlation between GEBVs obtained with the full inverse and GEBVs obtained with the APY inverse. Furthermore, for the simulated data we assessed the accuracy for validation animals as the correlation between their GEBVs and true breeding values (TBV). For the real pig data we assessed the accuracy for validation animals as the correlation between their GEBVs and phenotypes adjusted for the fixed effects in the model. We consider the accuracy with the full **G** inverse as the baseline.

To gain insight into where the chosen core animals are positioned with regards to each other, we have visualised the population structure of genotyped animals with dimension reduction techniques. We used the classical linear Principal Component Analysis (PCA) and the novel non-linear Uniform Manifold Approximation and Projection (UMAP; [27]). We obtained UMAP using the `umap R` package [28] with default parameters.

### Computations

Simulated cattle data was analysed within the `R` environment [24], including the construction of genomic relationship matrices and their inverses by direct or APY inversion, and by calling the `BLUPF90` software [29] to solve the mixed model equations. Due to the size, the real pig data was solved by preconditioned conjugate gradient algorithm as implemented in the `BLUP90IOD2` software [30] with convergence criterion set to 10^−12^ and executed on The University of Edinburgh High-Performance Computing environment (Edinburgh Compute and Data Facility; http://www.ecdf.ed.ac.uk). For the simulated and real data we assumed that variance components are known and hence were not estimated from the data. All figures were produced using `ggplot2` [31] and `VennDiagram` [32] `R` packages. The code for the simulation, as well as core subset construction and genetic evaluation, is available from https://github.com/HighlanderLab/ipocrnic_OptimisedCore4APY (in the code the standard conditional algorithm is named as “prior”, while the two computationally more expensive extensions are named “posterior max” and “posterior avg”).

## Results

The quality of APY approximation depends on the size of the core subset. For a given core subset size we can optimise its construction to ensure stability of GEBVs. We show this with correlations between the GEBVs obtained with the full inverse and GEBVs obtained with the APY inverses, as well as respective validation accuracies. By plotting the population structure with PCA and UMAP, we show how the conditional algorithm spreads core animals far away from each other in covariance sense. Instability of the random construction is illustrated by Venn diagrams of overlapping core animals between replicates. We present these results separately for the simulated cattle data and the real pig data.

### Simulated cattle data

For the simulated data, the number of core animals was 10, 50, 135, 326, 968, 1516, 2386, and 3129. These core subset sizes captured respectively 10, 30, 50, 70, 90, 95, 98, and 99% of variation in **G**.

Figure 1 shows correlations between GEBVs obtained with the full inverse and GEBVs obtained with the APY inverse, calculated either for the whole genotyped or validation population. The correlations increased with the increasing core subset for all four core subset constructions. The conditional construction had generally the largest correlation. Other constructions had comparable correlations. Correlation in the validation population was more sensitive to core subset size than the whole population. The correlations were greater than 0.99 when the captured proportion of variation in **G** was 98%. There was considerable variation in correlation between replicates with the random and weighted constructions, though this variation reduced as the core subset increased. With a large core subset (2386; 98% variation captured in **G**), the correlations between five consecutive replicates of 3000 validation animals were all 0.99, for random and weighted constructions.

**Figure 1.**
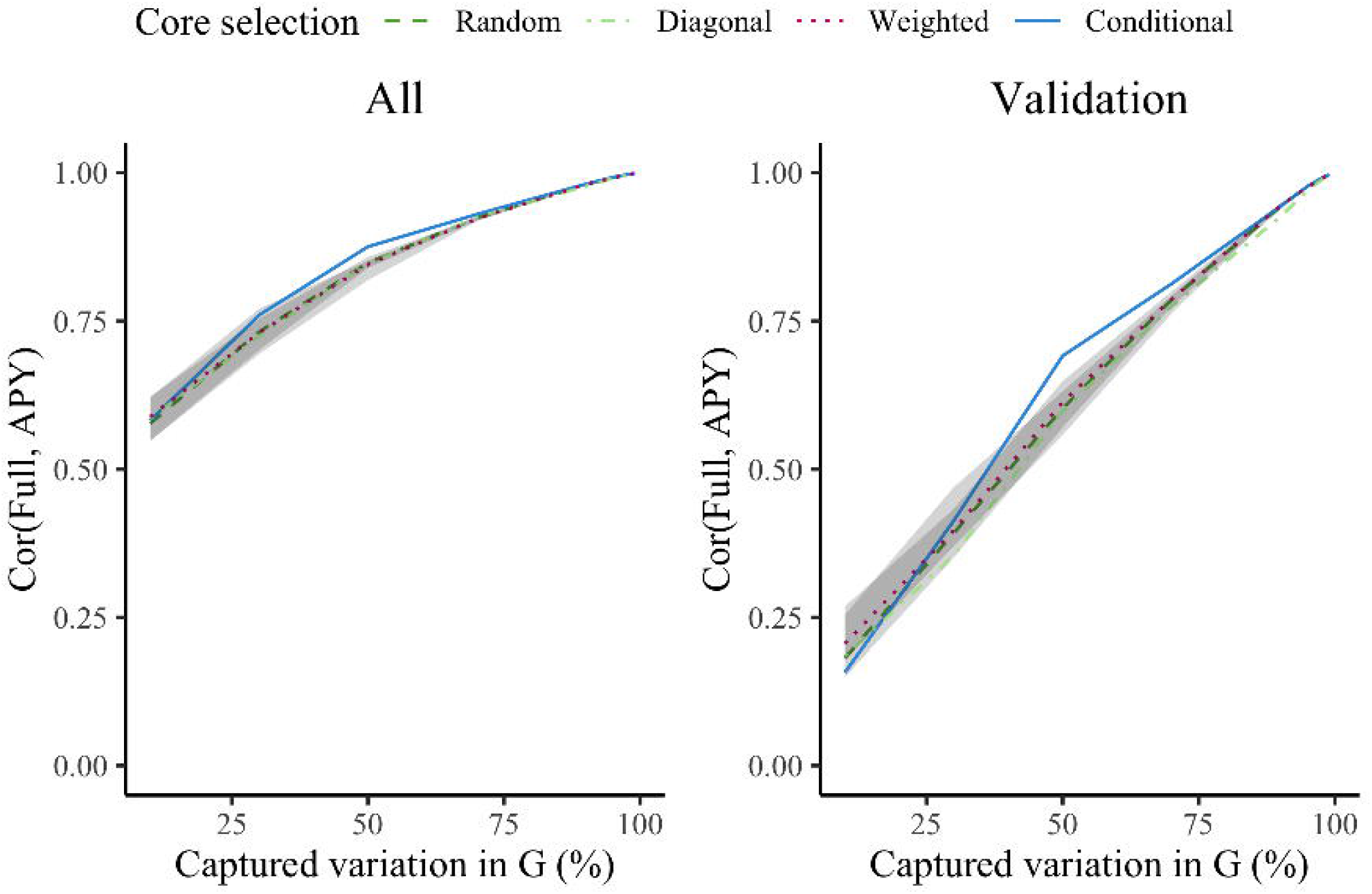
Correlations between GEBV_Full_ and GEBV_APY_ for all and validation animals in simulation. Genomic estimated breeding values (GEBV) were based on the full inverse (Full) or the Algorithm of Proven and Young inverse (APY) of the genomic relationship matrix (**G**). For APY, core subset was constructed at random (Random), based on the highest diagonal in **G** (Diagonal), combination of Random and Diagonal (Weighted), or conditional algorithm (Conditional). Random and weighted constructions show mean and 95% quantiles from five samples.

Figure 2 presents the accuracy of GEBVs for validation animals as a function of the percentage of captured variation in **G**. The accuracy with the full inverse was 0.74, which represents the baseline. The accuracy with the APY inverse increased from 0.10 to 0.75 as the core subset increased. There were only minimal differences between the four core subset constructions in accuracy trends, but we note the variation in accuracy with random constructions and that conditional construction generally had the largest correlation. The accuracy reached or even marginally surpassed (for about 0.001) the accuracy obtained with the full inverse, when 98% of the variation in **G** was captured.

**Figure 2.**
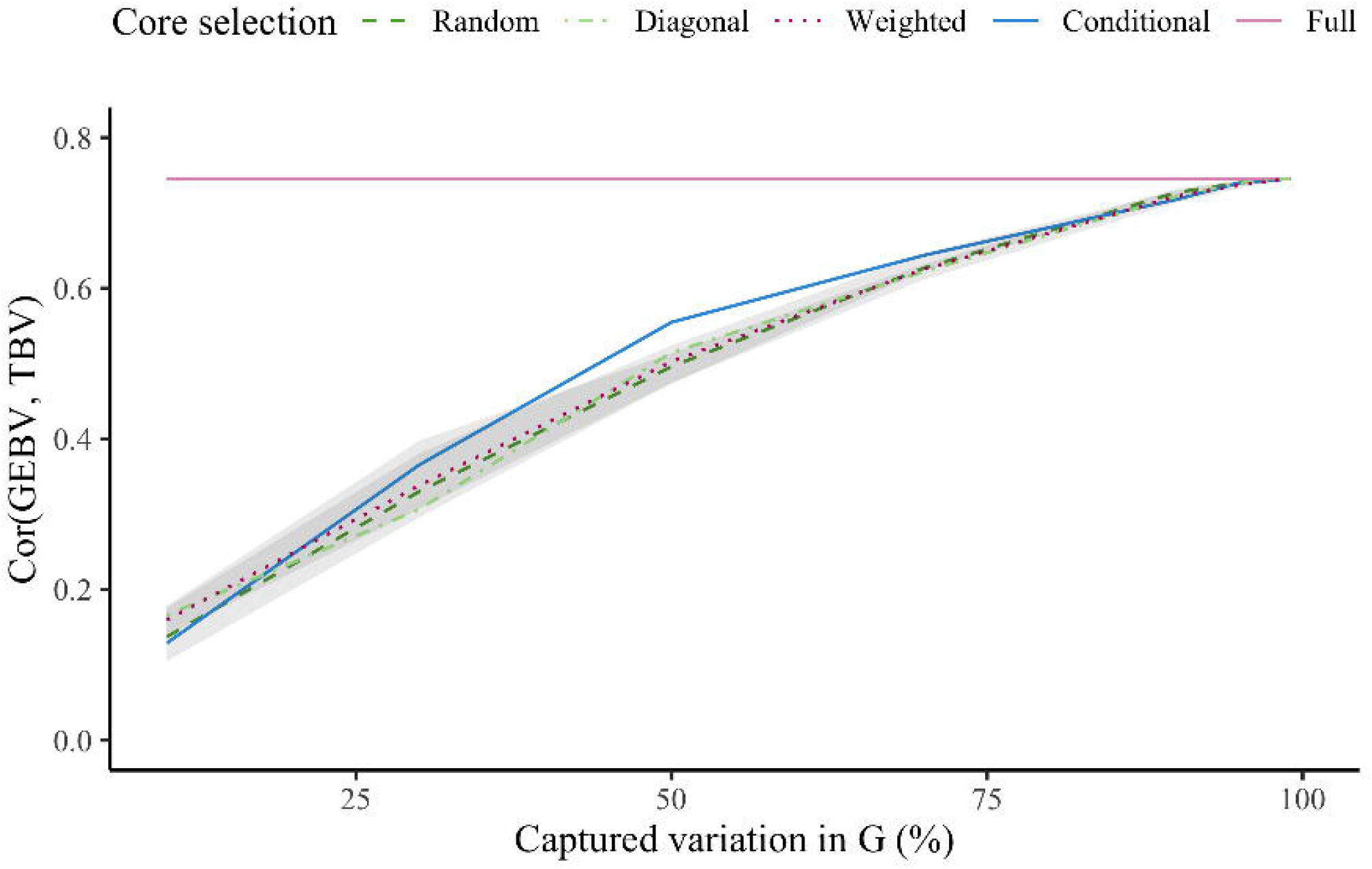
Accuracy for validation animals in simulation. Accuracy is the correlation between genomic estimates of breeding values (GEBV) and true breeding values (TBV) from the simulation. GEBV were based on the full inverse (Full) and the Algorithm of Proven and Young (APY) inverse of the genomic relationship matrix (**G**). For APY, core subset was constructed at random (Random), based on the highest diagonal in **G** (Diagonal), combination of Random and Diagonal (Weighted), or conditional algorithm (Conditional). Random and weighted constructions show mean and 95% quantiles from five samples.

To understand how the different core subset constructions may affect selection decisions, we compared the top 3% (45 out of 1500) of young males selected based GEBVs from the full inverse or the APY inverse. These selected young males were used as sires of the next generation so their selection is important. When the core subset captured 98% of variation in **G**, very similar animals were selected using GEBVs calculated with the full inverse compared to the APY inverse. Specifically, compared to the full inverse, the random construction correctly selected 37.6±2.3 of the 45 sires, the diagonal construction correctly selected 37/45±0.0, the weighted construction correctly selected 39.2±1.3, and the conditional construction correctly selected 39/45±0.0. Therefore, random construction had more variation than weighted construction, with respectively 35-40 correctly selected sires compared to 38-40. There is no variation in diagonal and conditional constructions by design.

Compared to the random construction, the conditional construction could be time consuming because the conditional algorithm is updating a large SNP genotype matrix **W**. To speedup this potential time bottleneck, we also ran the conditional algorithm on a reduced rank SNP genotype matrix obtained via **W**_*r*_ = **U**_*r*_**D**_*r*_, where **U**_*r*_**D**_*r*_ correspond to the first *r* principal components of **W** that captured 99% of the variation in **G**, that is 3130 of the 15,000 principal components. Using the reduced rank SNP genotype matrix **W**_*r*_ with conditional construction produced very similar accuracies of the GEBV as using the full SNP genotype matrix **W** (Additional File 1). However, there were some differences between the chosen core animals when using **W**_*r*_ or **W**, which increased proportionally with the number of core animals. For example, with 10 core animals the overlap was 90% (9/10), with 135 it was 86% (116/135), but with 2386 it was only 71% (1696/2386). The time required to choose core animals using reduced rank **W**_*r*_, obtain the APY inverse in `R`, read the inverse and solve the mixed model equations in `BLUPF90`, was reduced by more than half compared to using the full **W**.

Additional File 2 shows PCA and UMAP for the 15,000 animals in the simulation, with data points coloured by generation. While both methods captured general homogeneity of the simulated population, UMAP also revealed fine-scale population structure and more pronouncedly revealed change of variation between generations (Additional File 3). Namely, the projected points moved closer as the generation number increased. This trend suggests loss of genetic variation due to selection and the creation of distinct clusters of paternal half-sib families. For example, the 50 clusters identified by UMAP in the validation set (generation 20) correspond to exactly 50 paternal half-sib families (Additional File 2, 3).

With a clear population structure of paternal half-sib families in the validation population, we investigated how the conditional algorithm spreads core animals. In Figure 3 we visualise UMAP for the validation population and mark chosen core animals. With just a few exceptions, the conditional algorithm spread the core animals across the validation population by selecting a core animal from each half-sib family. By comparison, the random construction would randomly spread core animals across the validation population, each time differently.

**Figure 3.**
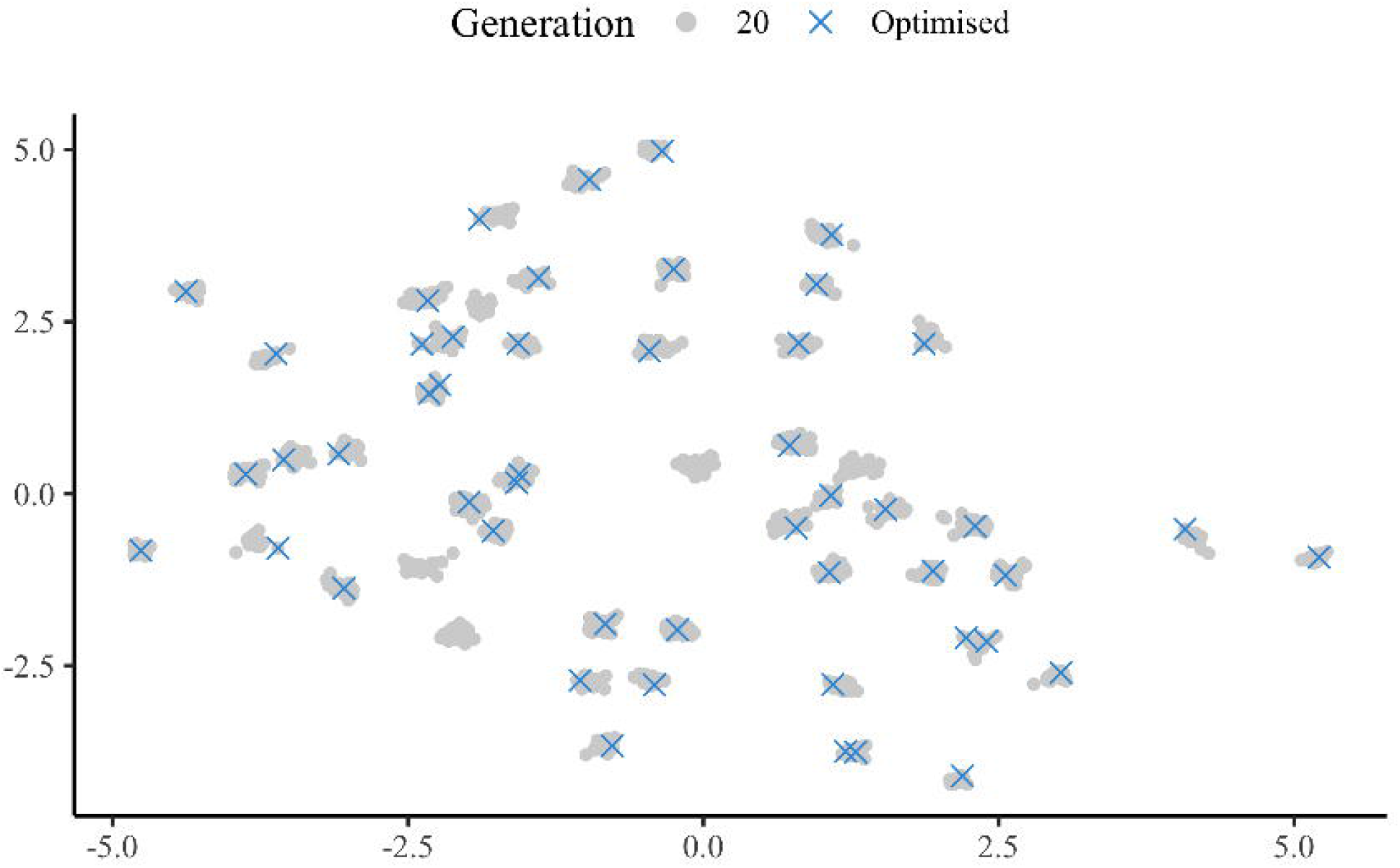
Spread of core animals in the last generation of simulation. Visualisation of the Uniform Manifold Approximation and Projection (UMAP) for animals (gray dots) in the last generation of simulation representing 50 paternal half-sib families. Overlaid are 50 core animals chosen by the conditional algorithm (blue crosses).

Additional File 4 shows UMAP for the 15,000 animals in simulation and 10 core animals with the four core subset constructions. Random constructions show core animals from five samples to indicate variability. This figure again demonstrates how the core animals are spaced across a population and how these core animals differ between independent constructions. For example, no core animals overlapped across all five samples, and on average only 15% of core animals overlapped between pairs of replicates. By design, there is no such variability in the diagonal and conditional constructions - core animals are always the same for a given core subset size and a given set of genotyped animals. Note that some chosen core animals are not displayed since they have been placed outside the chosen axis limits - into two paternal half-sib families in generation 16.

### Real pig data

For the real data, the number of core animals was 8, 60, 184, 485, 1658, 2926, 5546, and 8348. These core subset sizes captured respectively 10, 30, 50, 70, 90, 95, 98, and 99% of the variation in **G**.

The Venn diagram in Figure 4 shows the number of shared core animals between the four core subset constructions. The total number of core animals is 5546, which captured 98% of the variation in **G**. The most of core animals were shared between the diagonal and conditional constructions (34%, 1873/5546), while less than 1% (29/5546) were shared across all four core subset constructions. The random construction produced the most unique core subset, with 73% (4045/5546) of core animals not shared with any other construction. For the random and weighted constructions, the results were consistent across all replicates, but we shown only a single replicate for clarity. Furthermore, the Venn diagram in Additional File 5 shows the number of shared core animals between five replicates of the random construction. In particular, there were no core animals shared across all 5 replicates, but at least 392 shared between pairs of replicates. A similar result was observed for the simulated data (not shown).

**Figure 4.**
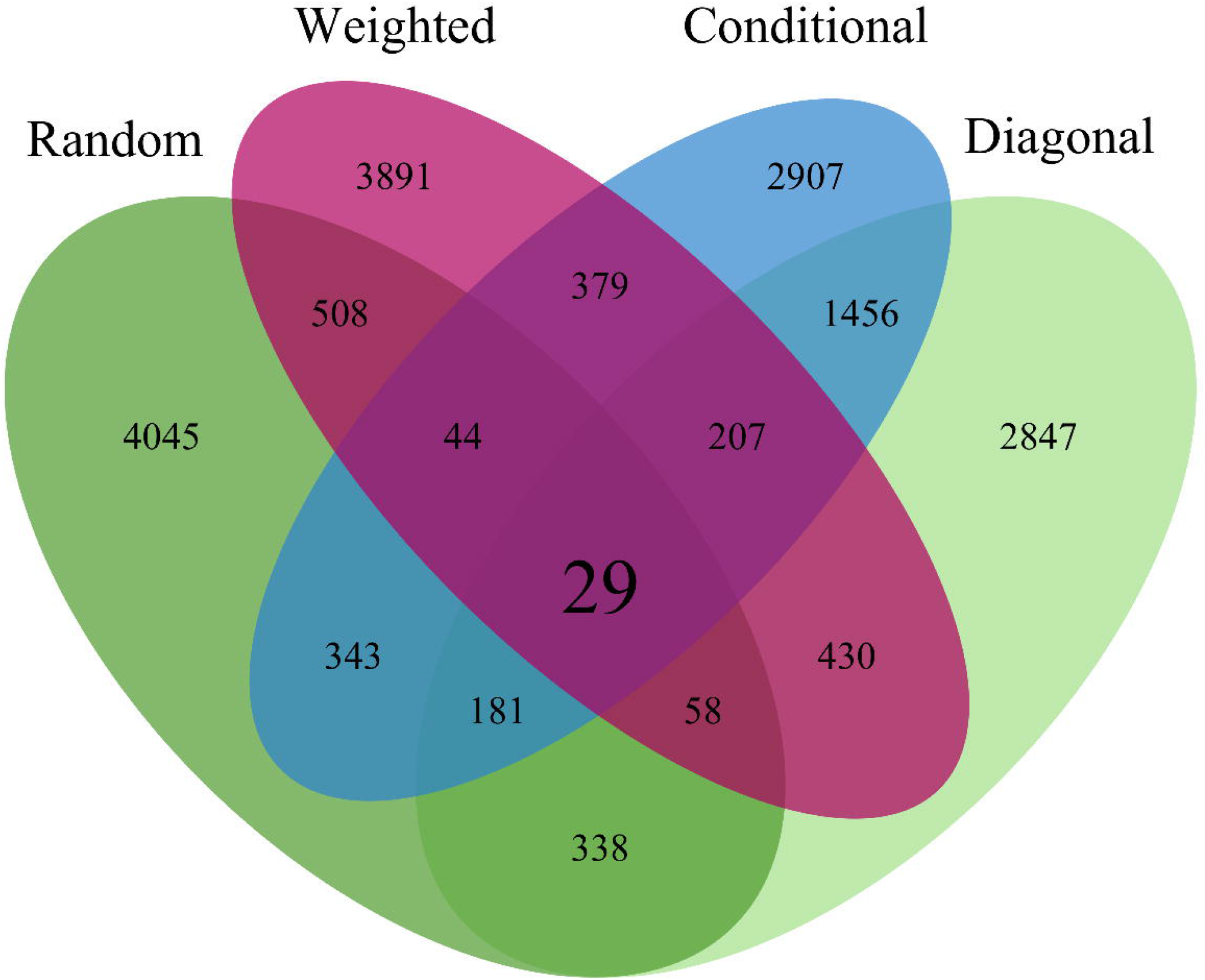
Venn diagram of core animals from different core subset constructions in pigs. Venn diagram is showing overlap for 5546 core animals between four core subset constructions. Core animals were selected at random (Random), based on the highest diagonal in in **G** matrix (Diagonal), combination of Random and Diagonal (Weighted), or conditional algorithm (Conditional). For Random and Weighted single replicate is shown.

Figure 5 shows that correlations between GEBVs obtained with the full and APY inverse in real data were almost a linear function of the percentage of variation captured with the APY approximation. All core subset constructions performed similarly well when the core subset was large, with correlations greater than 0.99 when 98% of the variation was captured in **G**. The difference between the four constructions was greater in the validation subset than in the whole genotyped population. The diagonal construction produced lower correlations than the other constructions for the full genotyped population, but not for the validation subset. Furthermore, the 95% quantiles for the random and weighted constructions were large when the number of core animals was low, especially for the validation subset. This indicates there was a large degree of variability between the five replicates of random and weighted construction. Conditional construction often had the highest correlation, particularly in the validation subset, but not always.

**Figure 5.**
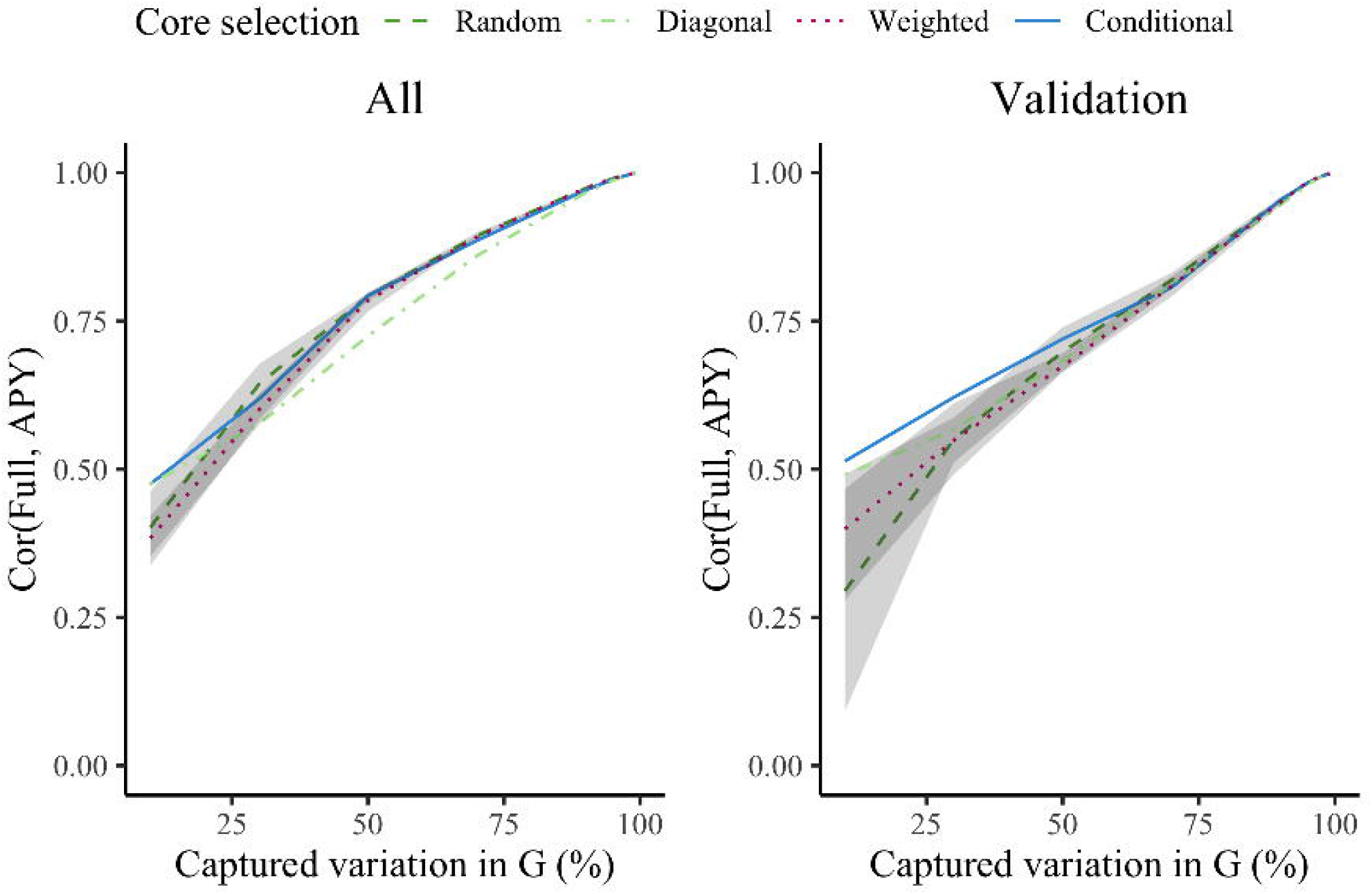
Correlations between GEBV_Full_ and GEBV_APY_ for all and validation pigs. Genomic estimated breeding values (GEBV) were based on the full inverse (Full) or the Algorithm of Proven and Young inverse (APY) of the genomic relationship matrix (**G**). For APY, core subset was constructed at random (Random), based on the highest diagonal in **G** (Diagonal), combination of Random and Diagonal (Weighted), or conditional algorithm (Conditional). Random and weighted constructions show mean and 95% quantiles from five samples.

Figure 6 shows the accuracy of GEBVs for validation animals relative to the percentage of variation captured in **G**. For example, the accuracy of GEBVs obtained with the full inverse was 0.31, and when 90% of the variance was captured in **G**, the random construction had accuracy of 0.28 (from 0.27 to 0.30), the diagonal construction 0.29, the weighted construction 0.27 (from 0.26 to 0.29) and the conditional construction 0.29. Again, the conditional construction generally had the highest accuracy. The accuracy of the random construction was higher than the conditional construction when more than 95% variance in **G** was captured, but note that the plotted difference is more pronounced by the rounding of accuracies to two decimal digits. The difference between the two constructions was always less than 0.005. The diagonal construction marginally surpassed accuracy of the full inverse when 99% of the variation in **G** was captured. In this sense, the conditional construction achieved the same satisfactory accuracy as the random construction when the number of core animals was large, but also achieved notable improvements over the random construction when the number of core animals was smaller. This improvement can be demonstrated by comparing accuracy from the conditional construction with the large variability in accuracy from the random and weighted constructions for a small number of core animals.

**Figure 6.**
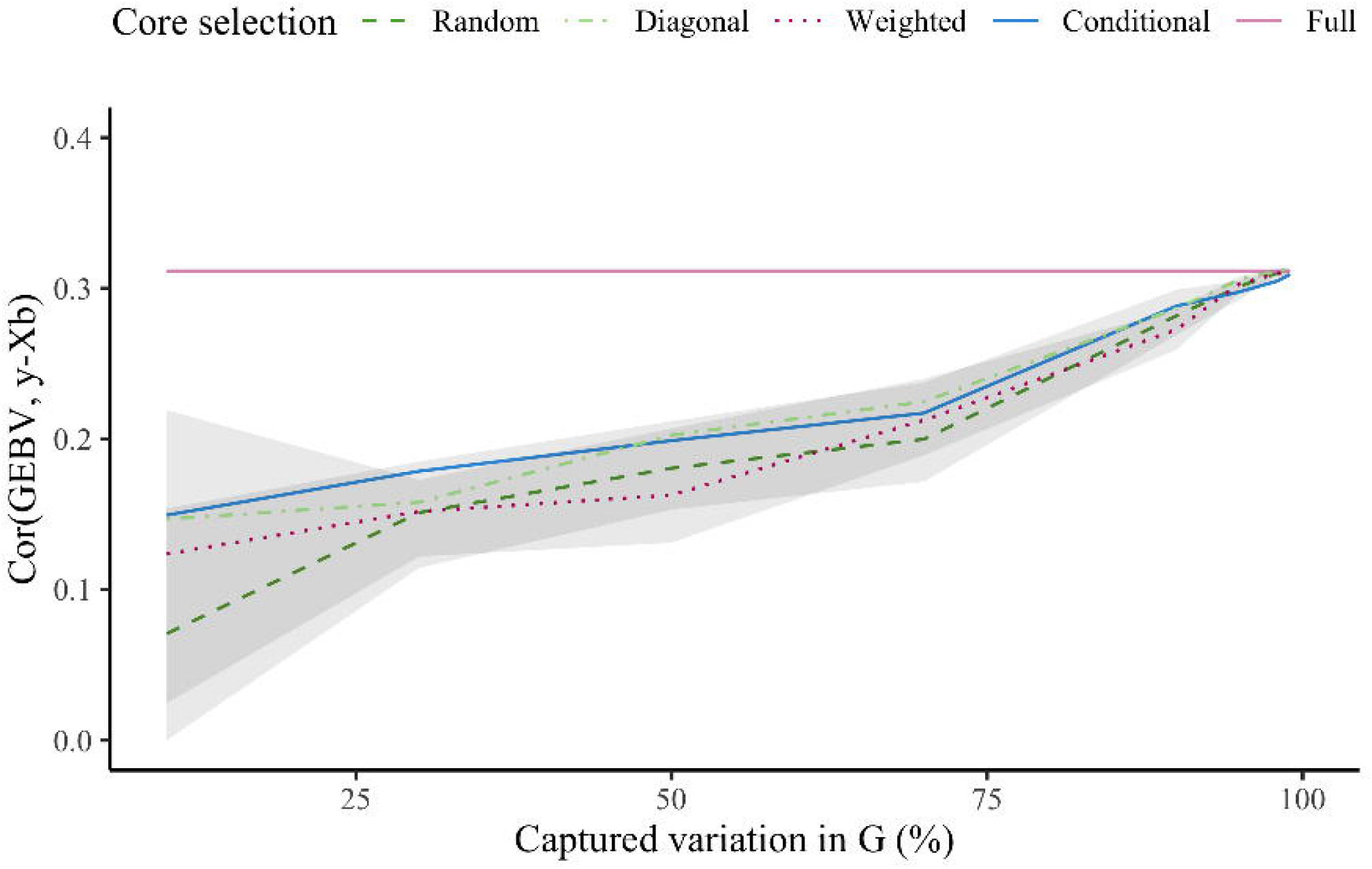
Predictive ability for the validation set in pigs. Accuracy is the correlation between genomic estimates of breeding values (GEBV) and phenotypes adjusted for the fixed effects (**y** − **Xb**). GEBV were based on the full inverse (Full) or the Algorithm of Proven and Young (APY) inverse of the genomic relationship matrix (**G**). For APY, core subset was constructed at random (Random), based on the highest diagonal in **G** (Diagonal), combination of Random and Diagonal (Weighted), or conditional algorithm (Conditional). Random and weighted constructions show mean and 95% quantiles from five samples.

We visualised structure of the pig population with PCA and UMAP as shown in Additional File 6. The PCA showed clear clusters of crossbreed animals (F1, BC1, and BC2) and purebred animals (L1). As expected, the backcross animals were positioned between the purebred and F1 animals. The UMAP also showed clusters of animals, but while the PCA showed L1 as a homogeneous population in first two dimensions, the UMAP revealed additional structure within L1 animals in first two dimensions. Closer inspection of the data showed that this additional structure corresponds to a time trend within a breeding programme (not shown).

Figure 7 illustrates how the conditional algorithm spreads animals, with an example of 60 core animals to facilitate visualisation. In this example, the conditional algorithm has chosen 14 BC1, 13 BC2, 1 F1, and 32 L1 animals. For comparison, one random sample included 11 BC1, 5 BC2, 2 F1, and 42 L1 animals.

**Figure 7.**
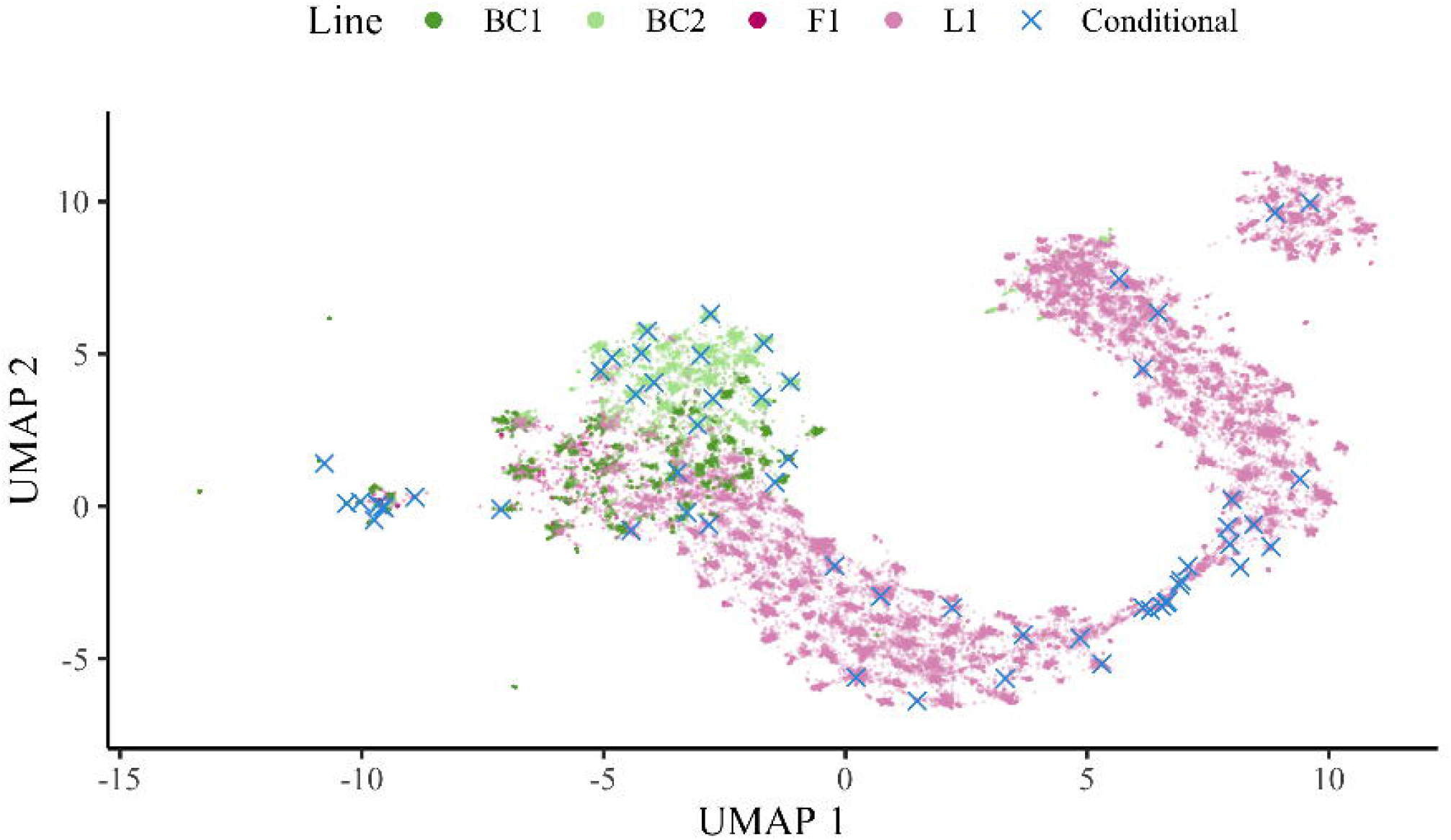
Visualisation of UMAP for genotyped pigs and an example of optimised core subset for 60 pigs. Animals chosen by the conditional algorithm (Conditional) are plotted on the Uniform Manifold Approximation and Projection (UMAP). Colours represent purebred (L1), crossbred (F1), and backcross (BC1, BC2) pigs.

## Discussion

The results show that core subset construction can be optimised and that such optimisations delivers accurate and stable GEBVs. Our results raise five points for discussion: (i) the overall performance of APY and the need for optimising the core subset; (ii) towards optimal core subset construction; (iii) expanding the core subset with new genotype data; (iv) limitations; and (v) other opportunities.

### The overall performance of APY and the need for optimising the core subset

Our results confirm that the size of the core subset is critical in APY and that robust results can be obtained with a large core subset. The results further show that given the size of the core subset, we can use the proposed conditional algorithm to achieve a good and repeatable spread of core animals across the domain of genotyped animals. We observed a similar increase in accuracy with the increasing core subset size in the simulated and real data. However, there was more variability in accuracy with the random construction in the real data than in the simulated data. Furthermore, the difference between core subset constructions was more apparent in the real data, especially with a small core subset. This was most likely due to the more complex structure in the real data, which shows the need for optimisation. Although the relationship between the size of the core subset and the accuracy of GEBVs is well described [17, 18, 20, 33], construction of the core subset is somewhat neglected, with the random construction being the most common. The proposed conditional construction contributes to addressing this situation. While it requires more time than the random construction, this additional time could be justified in several cases, specifically when the genotyped population has a complex structure, like multiple breeds or crossbred animals.

Ostersen *et al*. [13] and Bradford *et al*. [18] provided early indications that the choice of animals for the core subset is important in specific cases. For example, in the case where genotyped animals had an incomplete pedigree (with a variable number of generations), spreading the core animals across multiple generations maximised accuracy [18, 13]. As shown in our results, complex population structures, such as when the data includes multiple breeds or crossbred animals, necesitate some optimisation of the core subset. Mäntysaari *et al*. [8] mentioned this issue in their genetic evaluation involving 41 breeds, but did not provide a recommendation on how to optimise the core subset. Vandenplas *et al*. [34] simulated a three-way crossbreeding program, and observed that the best accuracy was achieved when the core animals were randomly chosen within each breed and crossbred population. This simulation was later corroborated with real data [14]. Vandenplas *et al*. [34] also suggested that statistically or numerically more “stable” core subsets might be needed for populations with a complex population structure, even if this increases computing cost. Their approach with the QR decomposition of SNP genotype matrix choose a core subset that improved convergence, but it did not improve accuracy. Nilforooshan and Lee [15] used the diagonal construction based on **G** as well as based on **A**. They found that the diagonal construction based on **G** gave similar correlation between GEBVs from the full inverse and APY inverse, when the core subset was large, but worse with smaller core subsets. We observed similar results with the real data. Recently, Cesarani *et al*. [35] used APY in a large-scale genomic evaluation including five breeds. They observed changes in the prediction accuracy with different choices of core animals. They concluded that the random choice of core animals impacts prediction accuracy because breeds with less genotyped animals are not well represented. To address this undersampling, they proposed increasing the core subset size and balancing the core subset size per breed. In their case this lead to the core (total number of genotyped animals) including 15k (3.4M) Holstein, 15k (400k) Jersey, 5k (9k) Ayshire, 5k (47k) Brown Swiss, and 5k (5k) Guernsey animals. Nevertheless their within-breed core subsets were still random. The conditional algorithm could optimise such a core subset because it works with SNP genotypes that capture both between and within breed variation.

### Towards optimal core subset construction

To our knowledge, there were no attempts to statistically optimise the APY core subset, except from the perspective of ensuring better convergence [34]. All other attempts largely modified the random approach by adding additional criteria, like spreading random choice within each generation, group of animals (e.g., bulls, cows, most inbreed, popularity, etc.), or breeds [2, 13, 18, 34, 15, 20]. Our conditional core subset construction is following ideas from the sequential sampling design, in which sense it uses variance of SNP genotypes for each individual (that is, diagonal of the **G**) conditional on the current core subset. Using the conditional variance it chooses core animals far away from each other in covariance sense. This in turn spreads core animals across the domain of genotyped animals and equivalently the domain of collected SNP genotype data.

To gain insight about the spread of core animals across population we used UMAP. In simulated data, UMAP clearly revealed generations and half-sib families within generation. It also revealed that the conditional construction commonly chose one core animal per half-sib family. On the other hand, the random construction samples core animals at random and can sub-optimally cover the domain of genotyped animals. In real data, UMAP also clearly revealed population structure across breeds, time, and between family variation. Visualisation of the chosen core animals with the conditional construction in real data again showed how using conditional variance enables spread across the domain of genotyped animals with extensive population structure. Of note, UMAP on the real data produced some very large outliers, which we have omitted to improve visualisation (Additional File 7).

The results show how the conditional construction mitigates fluctuations in accuracy present with the random construction on smaller core subsets. Venn diagrams showed striking differences between different core subset constructions, but even more importantly between replicates of the random construction. For example, in our simulated and real data, the random construction had no overlap across five replicates. As expected, fluctuations in accuracy were larger with smaller core subsets and smaller with larger core subsets. While large core subsets are the norm and easy to accommodate on modern compute infrastructure, the conditional algorithm can be useful to reduce even the smallest fluctuations in accuracy. For example, in simulation we saw variation in selected sires even when the core subset captured most of variation in **G**.

It is important to point out that the conditional algorithm does not provide the globally optimal core subset. This was clearly seen in the results, where sometimes other core subset constructions had marginally higher accuracy. Namely, the conditional algorithm starts with one core animal and then sequentially grows the core subset in a repeatable manner by choosing an animal whose SNP genotypes are the least captured by the current core subset (that is, the animal has the largest conditional variance). But, a different start can produce a different core subset. For consistency, we have started with the animal with the largest diagonal value in **G**, hence the algorithm started on the farthest edge of the domain of genotyped animals. Alternatively, we could have started with the animal closest to the centroid of SNP genotypes, hence the algorithm would have started in the centre of the domain. While achieving the global optimality is hard, we attempted to improve the optimisation by sequentially choosing core animals that would maximise captured variation across the domain (that is, minimising conditional variance in the next iteration). However, such an algorithm was impractically slow. Further work is required to develop more optimal algorithms for core subset construction.

### Expanding the core subset with new genotype data

Optimising expansion of the APY core subset with the arrival of new genotype data has not been addressed in the literature although several empirical studies have been done [36, 37, 38]. When new genotype data arrives, we can either construct the core subset anew or expand the existing core subset. The proposed conditional algorithm can expand an existing core subset because it can take as an input original or conditional SNP genotype matrix. To expand the core subset with new genotype data we have to (1) condition the SNP genotype matrix of new animals on the current core subset, (2) combine the conditional SNP genotype matrices of old and new animals, and (3) run the conditional algorithm with the combined conditional SNP genotype matrix as an input. Conditioning SNP genotype matrix of new animals on the current core subset can be done efficiently with the Algorithm 3, where we combine SNP genotype matrices of core and new animals and sequentially condition the combined SNP genotype matrix on each core animal (line 5). We provide an implementation of this algorithm in our code. Our tests showed that running combined optimisation (Algorithm 2 on combined SNP genotype matrix) or expanding (Algorithm 2 on old SNP genotype matrix, Algorithm 3 on core and new SNP genotype matrix, and Algorithm 2 on conditional SNP genotype matrix) gave the same core subset.

#### Algorithm 3

Condition the SNP genotype matrix of new animals on a core subset

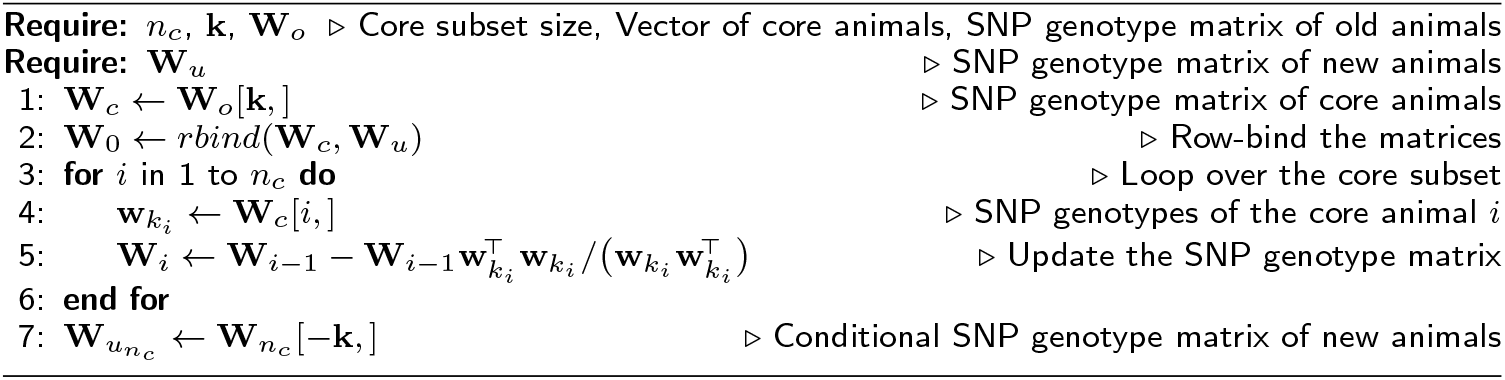

The described approach to expand the core subset assumes that the same allele frequencies are used in centering SNP genotype matrix of old and new animals. Ideally, we would use combined allele frequencies from old and new animals. However, the current core subset has been selected based on old allele frequencies. Hence, the same allele frequencies have to be used for new animals too. This is not ideal since some routine genetic evaluations continually update allele frequencies, though changes between genetic evaluations are likely small, even in breeding programmes. Some genetic evaluations use base population allele frequencies, which are expected to change even less.

Garcia *et al*. [37] tested the stability of APY-based indirect predictions when large numbers of genotyped animals are added to the data. They concluded that as long as the number of core animals is big enough, APY-based indirect predictions are stable, irrespective of core subset definition and addition of a large number of new genotypes. Similarly, Hidalgo *et al*. [38] examined the impact of adding new data with or without updating the core subset. They found that changes in GEBVs only slightly increased when the core was updated, compared to using the fixed core in the same period of time. Therefore, in line with the recommendations in Misztal *et al*. [19], Hidalgo *et al*. [38] recommend usage of the same core subset for a longer period of time (for example, one year), and update it when a significant amount of new data is generated. This update can be optimised with the conditional algorithm.

### Limitations

The main limitation of the conditional core subset construction is additional computation compared to the random construction. We see two options here. The first option is to work with a rank reduced SNP genotype matrix as we showed with the simulated data. There, iterating over the reduced rank SNP genotype matrix halved the time with no loss in accuracy, though core subsets were somewhat different. Therefore, while iterating over a rank reduced SNP genotype matrix saves time, it potentially adds another complexity, particularly when we also have to consider expansion with new data. The second option is to run the sequential algorithm for groups of animals by selecting them in the “selector” vector **e** instead of one animal. This change will likely produce a sub-optimal result and more research is needed on balancing compute time versus accuracy and stability of the APY approximation.

### Other opportunities

This study enables opportunities for further research in relation to at least four other study areas. First, the idea of using conditional variance for choosing core animals in APY is in fact identical to choosing animals for high-density genotyping or sequencing given a pedigree or low-density genomic relationship matrix [39]. Second, choosing core individuals that capture most of the genetic diversity is relevant also to designing and managing genebanks [40, 41]. Third, the conditional algorithm could be relevant to choosing a diverse subset for phenotyping where phenotyping is limited due to costs or other constraints [42, 43]. Fourth, inducing sparsity in dense inverse covariance (precision) matrices is an important topic in all areas that use Gaussian processes [44, 45]. One recent approach called Nearest Neighbour Gaussian Process (NNGP) gained popularity for large-scale applications in spatial statistics [46]. In genetics, NNGP would define a core subset for each animal, which is similar to the *Application 2* of Faux *et al*. [47] and ancestral regression Cantet *et al*. [48](when core subset represents parents and grand-parents). Extension of these concepts could further optimise APY for multi-breed applications. For example, linking the non-core animals of one breed only to the core animals of that breed, possibly also limit linking across too many generations to limit the contribution of older animals to recent generations.

## Conclusions

In summary, we have confirmed that accuracy of genomic evaluation with APY depends on the size of the core subset. For a given size of the core subset we can optimise the core subset with the proposed conditional algorithm. This algorithm achieves a good and repeatable spread of core animals across the domain of genotyped animals.

## Acknowledgements

Authors kindly thank Ignacy Misztal (University of Georgia, USA) for providing access to BLUP90IOD2 software. This work has made use of the resources provided by the Edinburgh Compute and Data Facility http://www.ecdf.ed.ac.uk/. Authors acknowledge valuable comments from Jana Obšteter (The Agricultural Institute of Slovenia, SI), Gabriela Mafra Fortuna (The University of Edinburgh, UK), and Thiago de Paula Oliveira (The University of Edinburgh, UK).

## Funding

GG, FL, and IP acknowledge funding from the Centre for Statistics (The University of Edinburgh). GG and IP acknowledge the BBSRC Institute Strategic Programme funding to The Roslin Institute (BBS/E/D/30002275). For the purpose of open access, the author has applied a CC BY public copyright licence to any Author Accepted Manuscript version arising from this submission.

## Availability of data and materials

R code for simulation and core subset construction are publicly available at https://github.com/HighlanderLab/ipocrnic_OptimisedCore4APY. The real pig data analysed in this study is owned by Genus PIC and is not publicly available.

## Ethics approval and consent to participate

Not applicable.

## Competing interests

The authors declare that they have no competing interests.

## Consent for publication

Not applicable.

## Authors’ contributions

IP designed the study, performed the analyses, and drafted the manuscript; FL conceptualised the conditional algorithm and contributed to the interpretation of results and discussion; WOH contributed to the design of the study and interpretation of results; DT contributed to the interpretation of results and discussion; GG initiated and supervised the study, helped with the design of the study, computations, interpretation of results and discussion. All authors read and approved the final manuscript.

## Additional Files

Additional file 1 — Accuracy of optimised core selection approach in simulation with rank reduction.

Here we show accuracy as the correlation between genomic estimates of breeding values (GEBV) and true breeding values in simulation. For he Algorithm of Proven and Young (APY), core subset was optimised on either full or rank reduced **W** matrix.

Additional file 2 — Visualisation of PCA and UMAP for genotyped animals in simulation.

Projection of genomic relationships into first two dimensions was done with Principal Components Analysis (PCA) or with Uniform Manifold Approximation and Projection (UMAP). The percentage of variation captured by each principal component is shown in parentheses. Colours represent five different generations of genotyped animals in the simulation.

Additional file 3 — Visualisation of UMAP over generations in simulation.

Here we visualise Uniform Manifold Approximation and Projection (UMAP) for genotyped animals in each of the five generations (16, 17, 18, 19, and 20) in the simulation.

Additional file 4 — Spread of core animals with different core subset constructions in simulation.

Ten core animals, from each of four core subset constructions, are plotted on the Uniform Manifold Approximation and Projection (UMAP). Core animals are either selected at random (Random), based on the highest diagonal in **G** (Diagonal), combination of Random and Diagonal (Weighted), or conditional algorithm (Conditional). Random and Weighted show core animals from five samples (1-5), Diagonal and Conditional from one sample (1), while non-core animals are shown as zero (0). “Missing” core animals were outside the chosen axis limits and are not displayed for the clarity of presentation.

Additional file 5 — Venn diagram of core animals from five random samples in pigs.

Venn diagram is showing overlap for 5546 core animals between five (1-5) random samples.

Additional file 6 — Visualisation of PCA and UMAP for genotyped pigs.

Projection of genomic relationships into first two dimensions was done with Principal Components Analysis (PCA) or with Uniform Manifold Approximation and Projection (UMAP). The percentage of variation captured by each principal component is shown in parentheses. Colours represent purebred (L1), crossbred (F1), and backcross (BC1, BC2) pigs.

Additional file 7— Visualisation of UMAP before and after cleaning in real data

Here we visualise Uniform Manifold Approximation and Projection (UMAP) for genotyped purebred (L1), crossbred (F1), and backcross (BC1, BC2) pigs, before (a) and after (b) removing 378 (*<* 1%) pigs from the final UMAP plot for the sake of clarity.

## Notes

### Competing Interest Statement

The authors have declared no competing interest.

