## Additional File 1 for "Optimisation of the core subset for the APY approximation of genomic relationships"

| Number of core animals | Percentage of variation explained in G | Accuracy with full W | Accuracy with rank reduced W |
|---|---|---|---|
| **10** | 10 | 0.13 | 0.13 |
| **50** | 30 | 0.37 | 0.36 |
| **135** | 50 | 0.55 | 0.53 |
| **326** | 70 | 0.64 | 0.66 |
| **968** | 90 | 0.72 | 0.73 |
| **1516** | 95 | 0.74 | 0.74 |
| **2386** | 98 | 0.74 | 0.75 |
