## Supplementary figures and images for "Optimisation of the core subset for the APY approximation of genomic relationships"

### Additional File 2

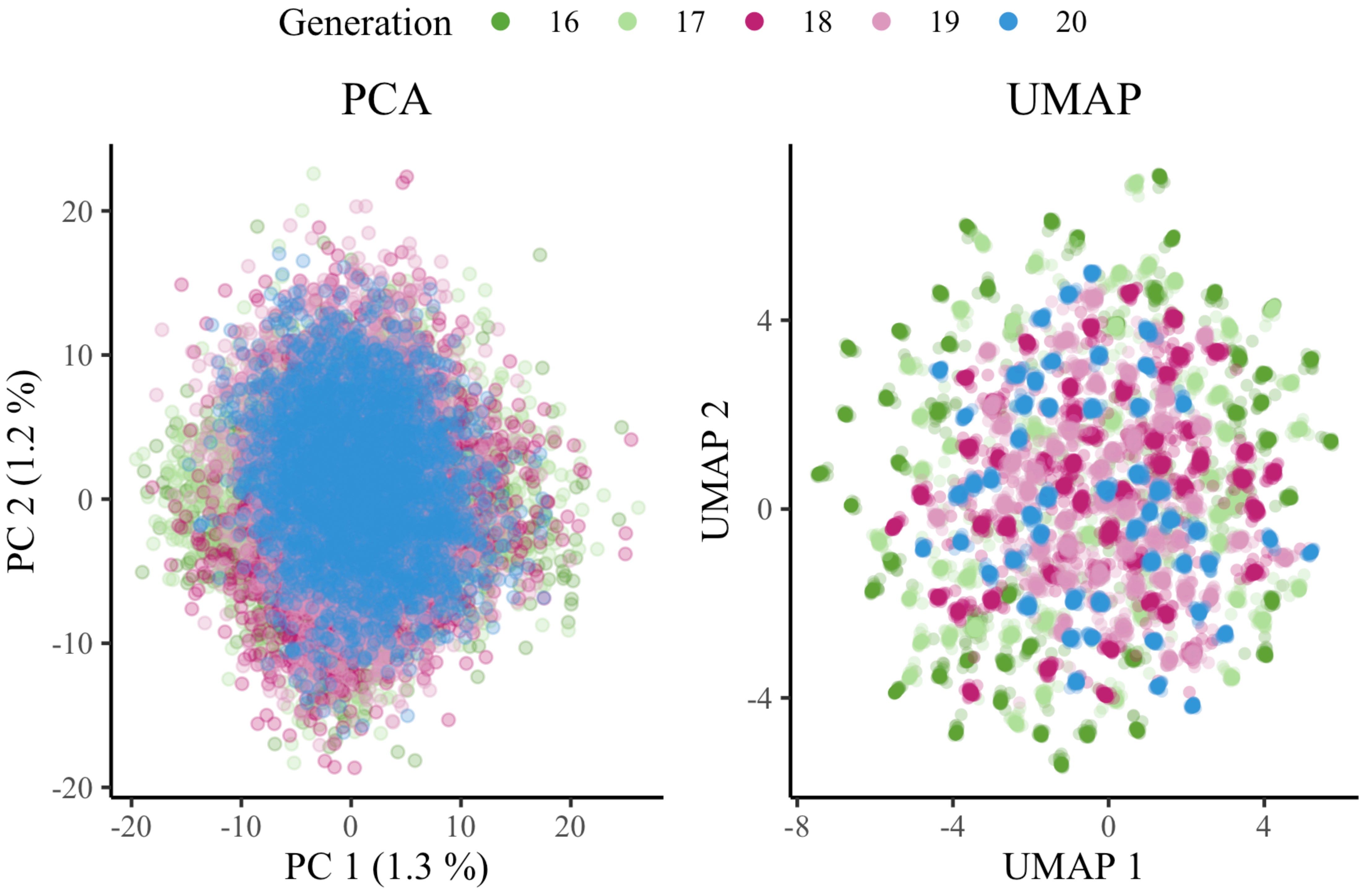

### Additional File 3

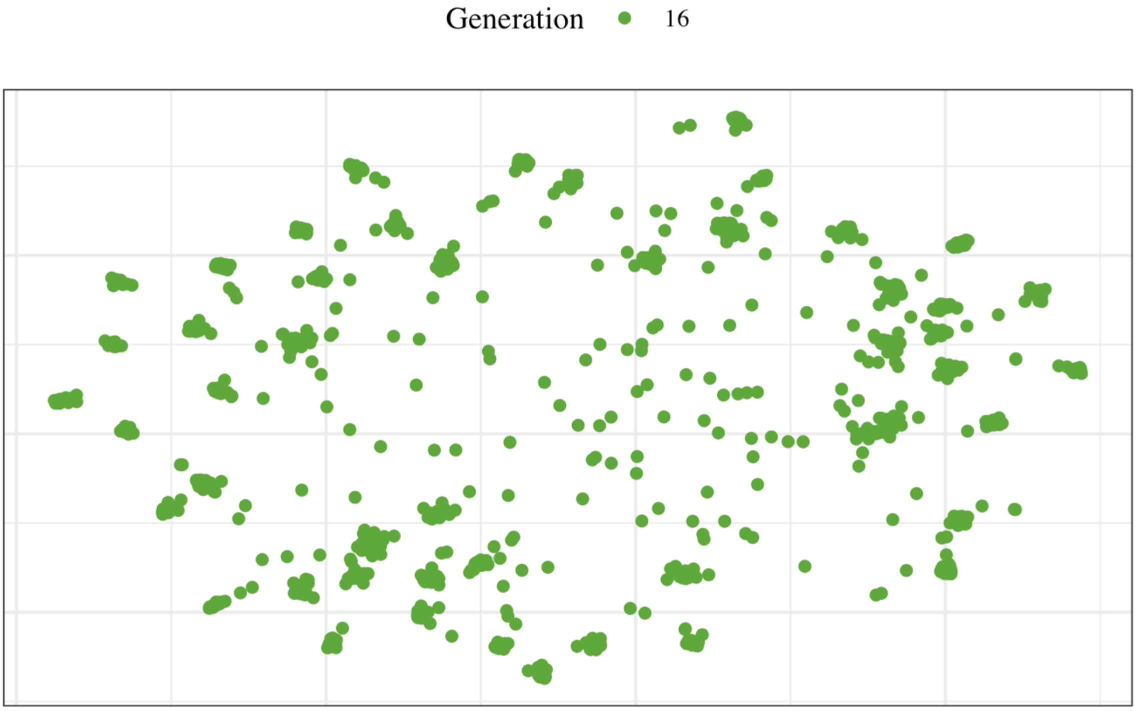

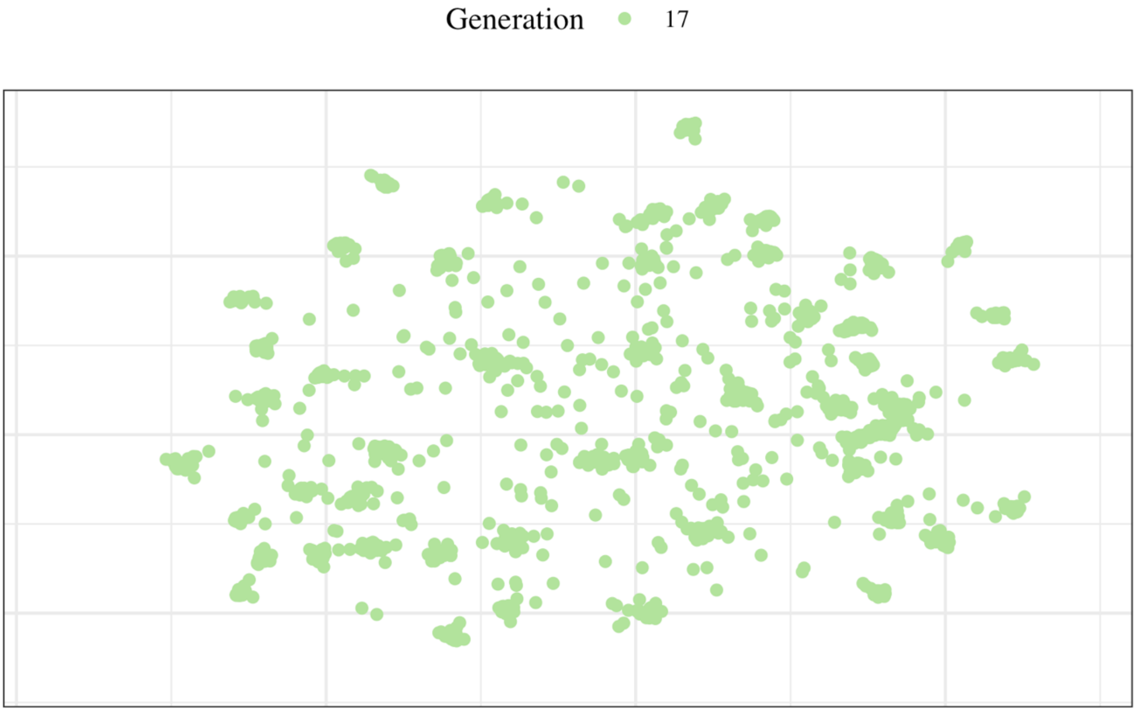

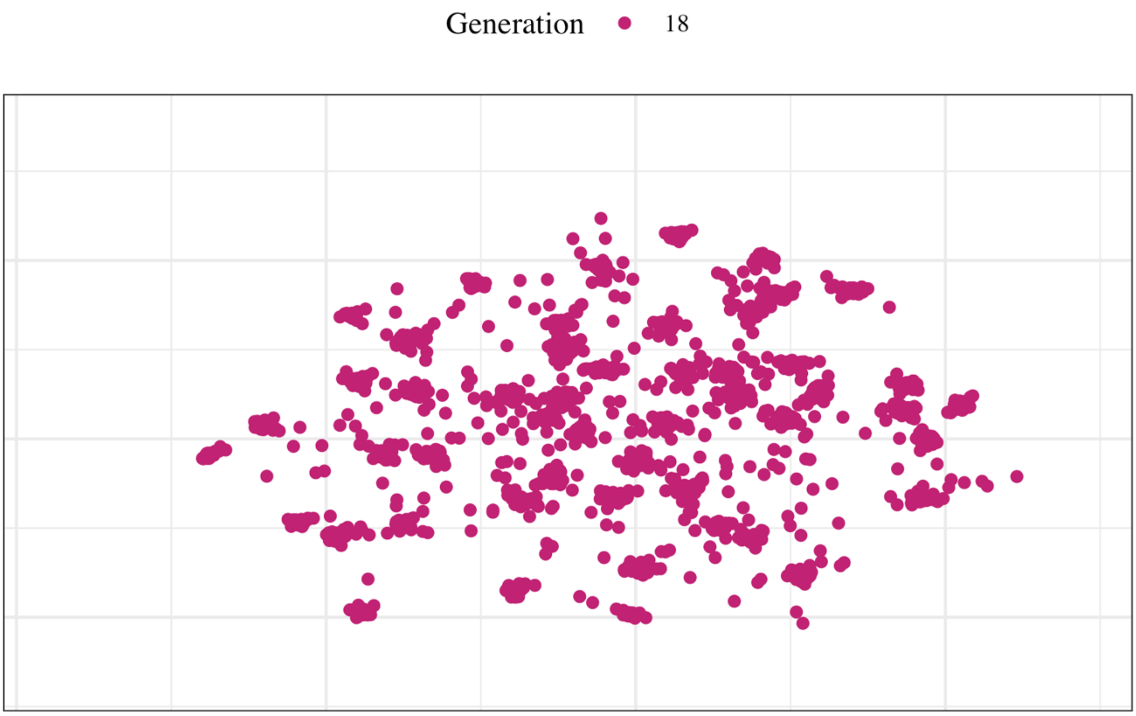

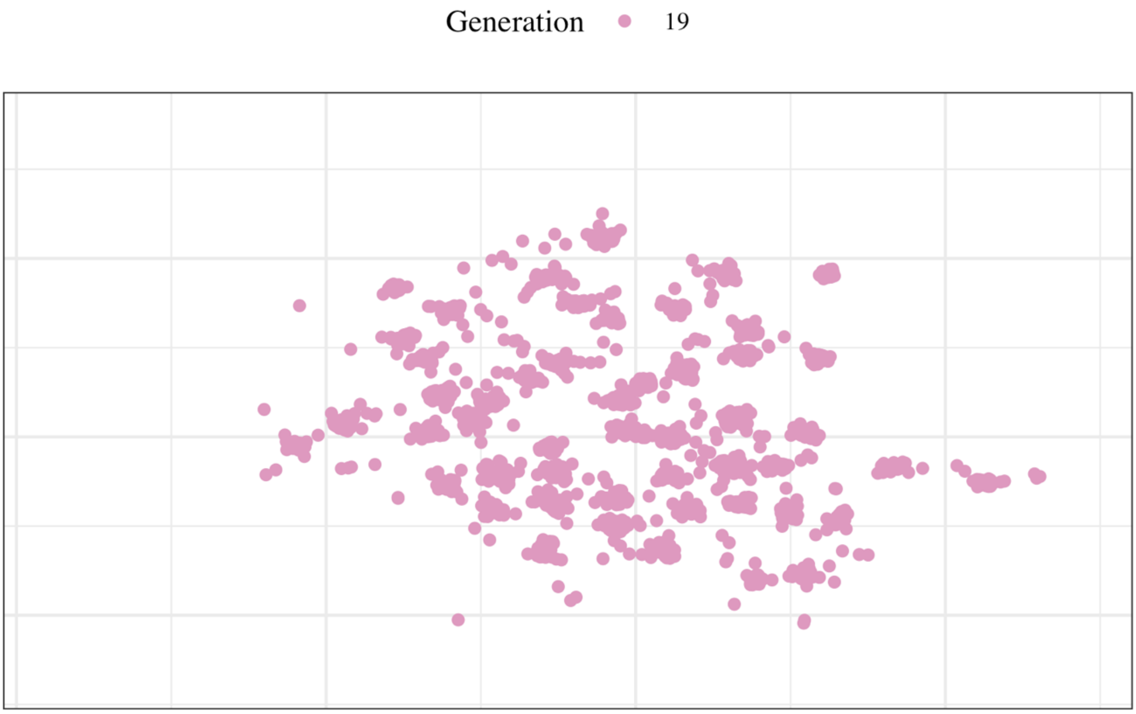

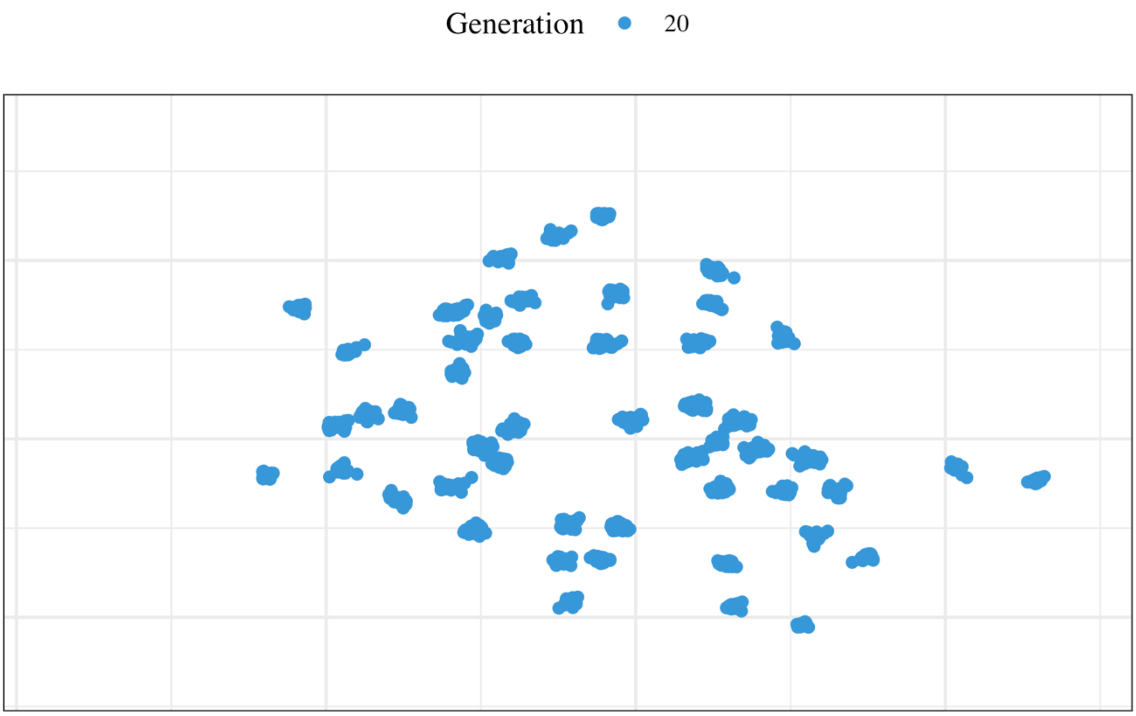

### Additional File 4

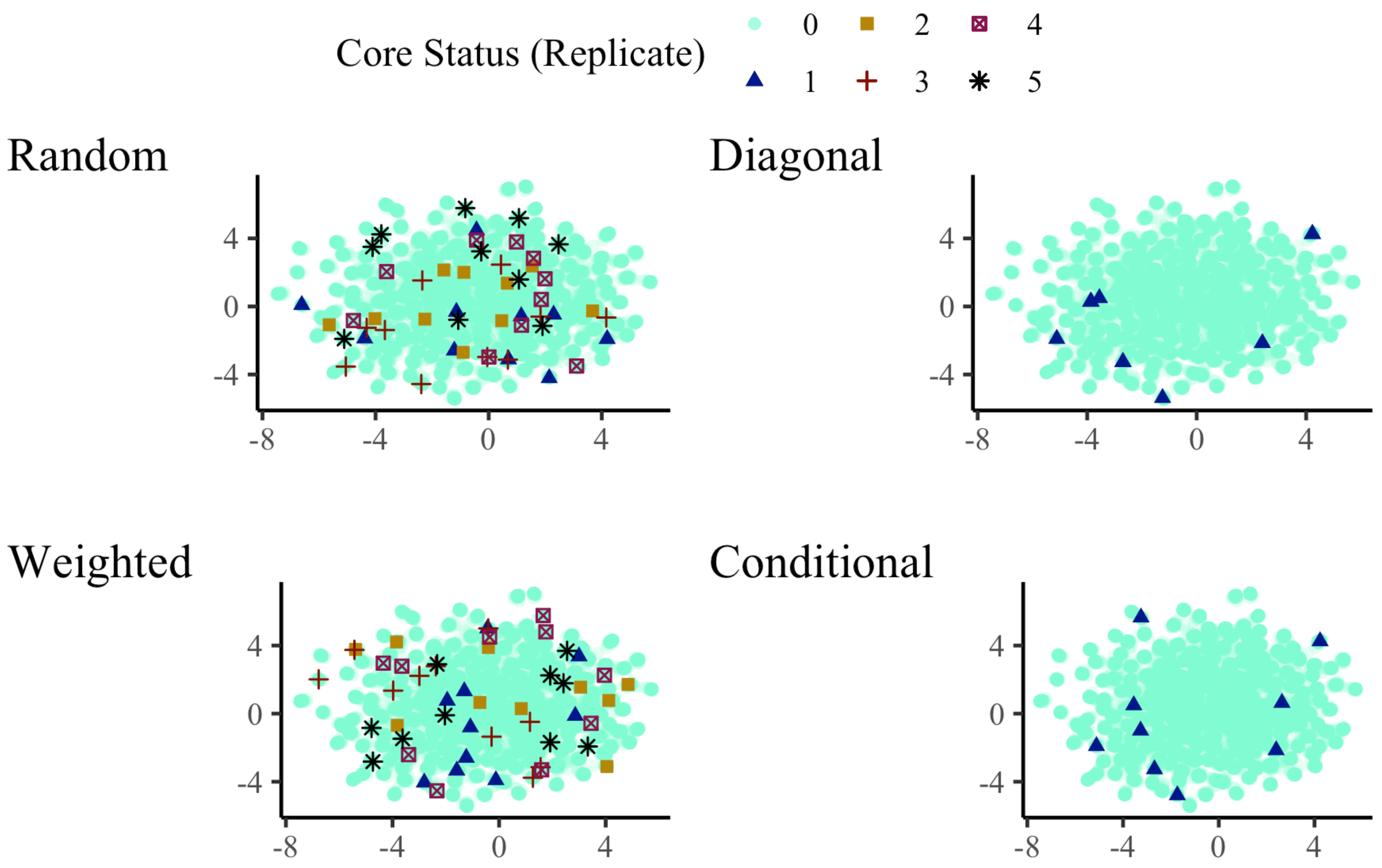

### Additional File 5

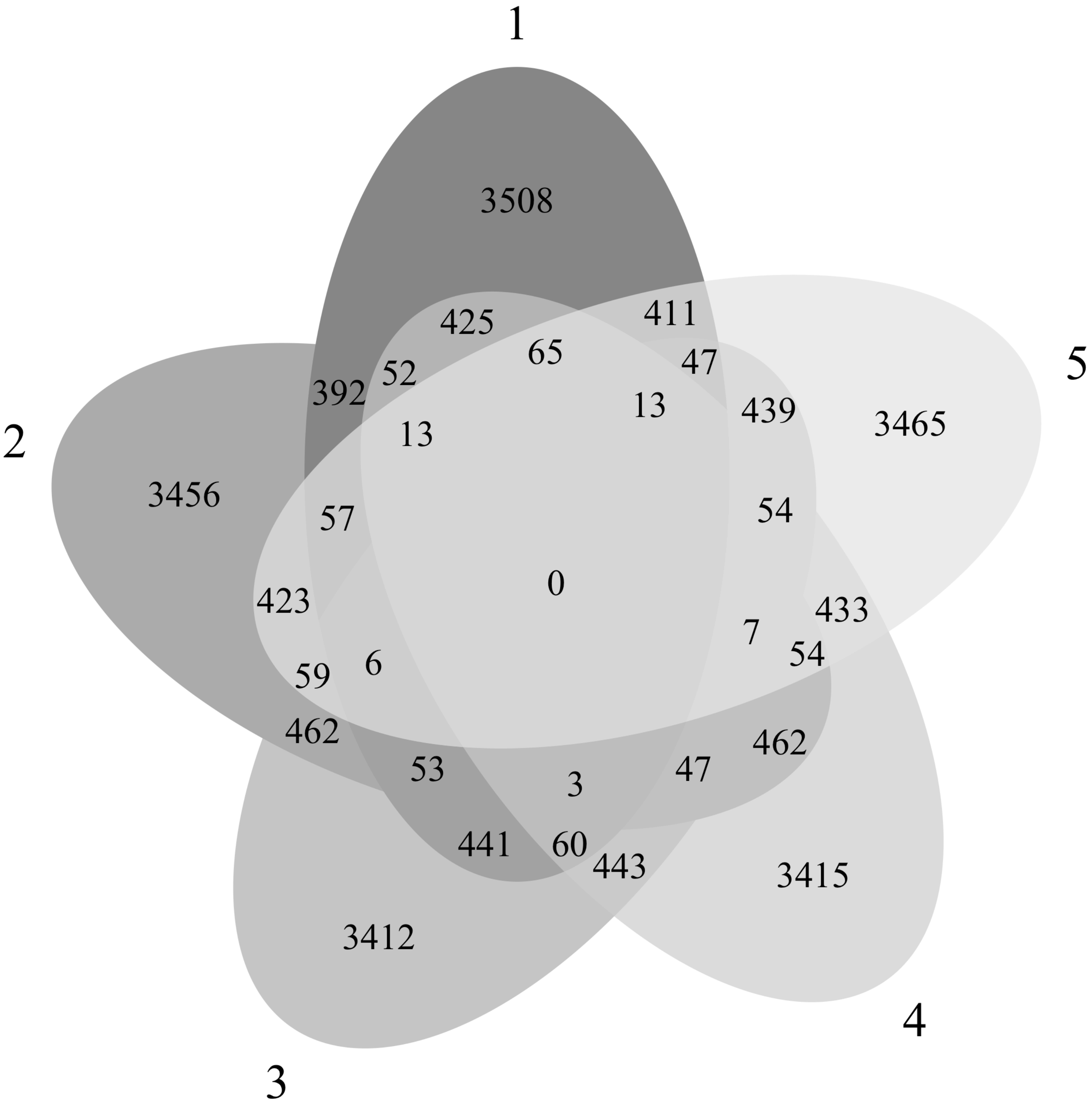

### Additional File 6

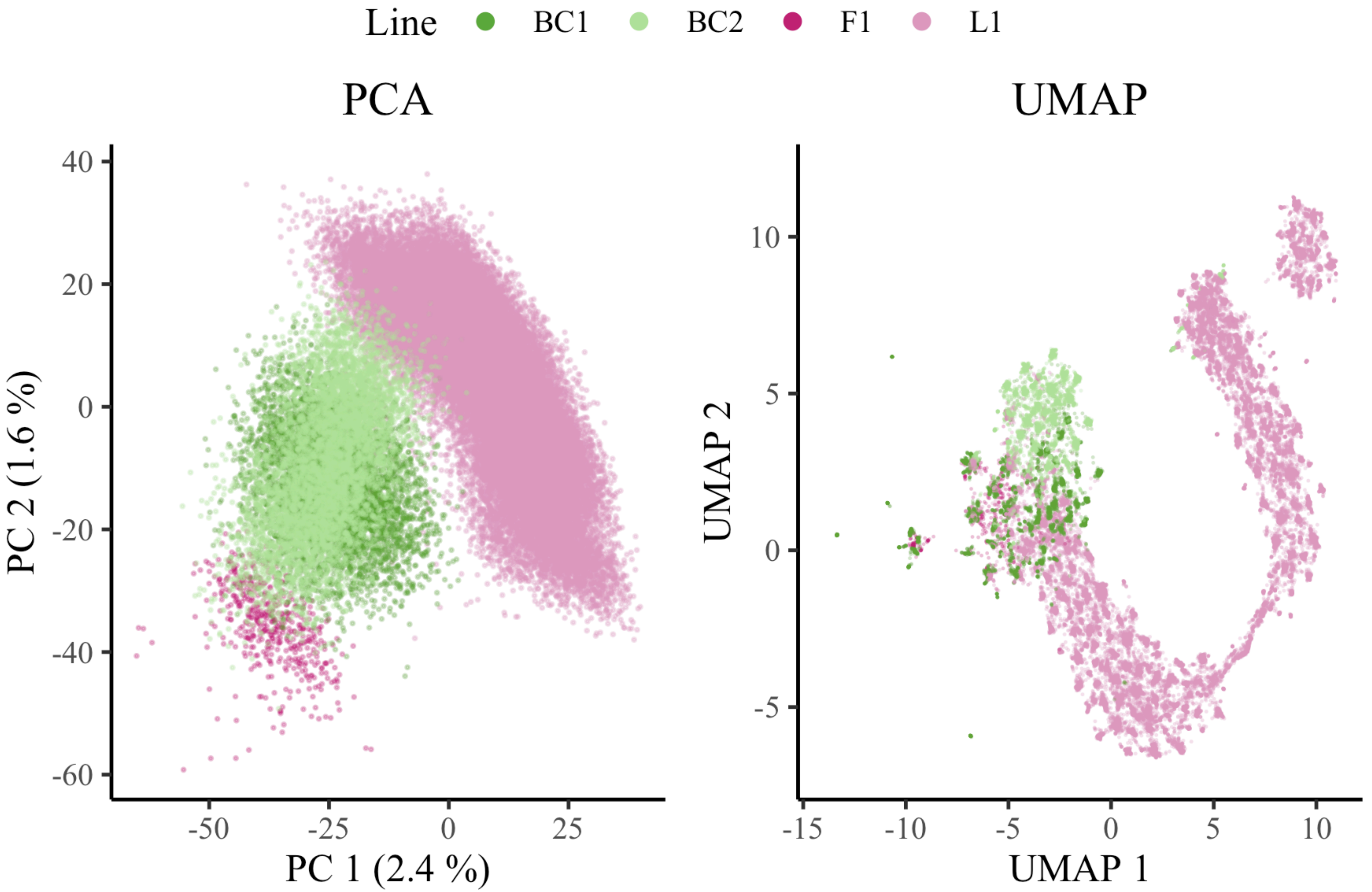

### Additional File 7

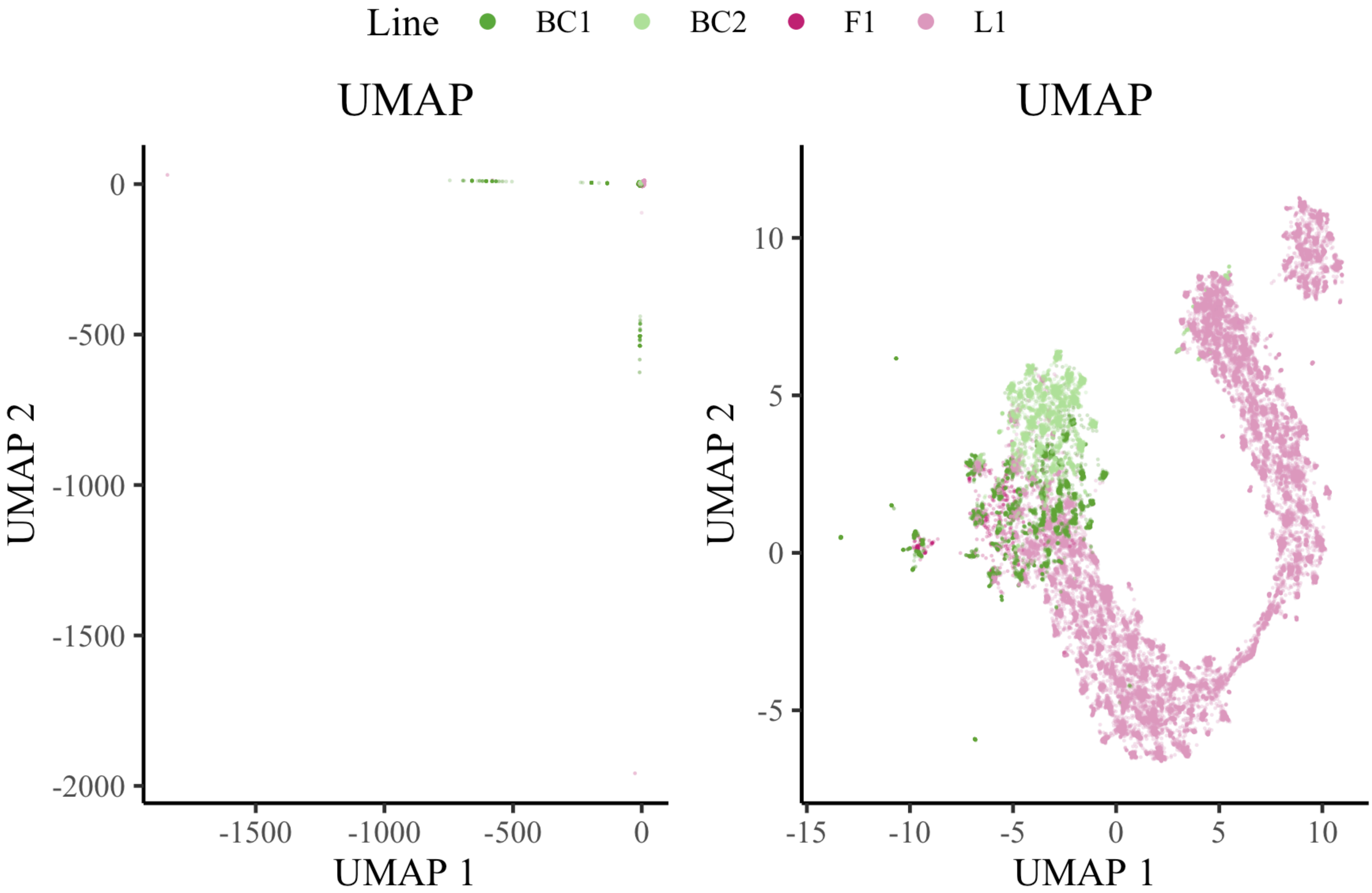
